# Using fisheries risk assessment to inform precautionary and collaborative management in a declining coho salmon fishery

**DOI:** 10.1101/2025.02.26.640431

**Authors:** Kyle L. Wilson, William I. Atlas, Charlotte K. Whitney, Mike Reid, Christina N. Service, Brendan M. Connors, Megan S. Adams

## Abstract

The conservation and management of Pacific salmon in Canada faces an uncertain future. While fisheries policies, like Canada’s Wild Salmon Policy, increasingly emphasize conservation, salmon continue to decline due to cumulative pressures of climate change, habitat loss, and overfishing, requiring precautionary management. We quantified population dynamics for 52 coho salmon populations along the North and Central Coast of British Columbia since 1980 to determine population status, assess risks posed by a mixed-stock fishery spanning U.S., Canadian, and Indigenous jurisdictions, and implemented forward simulations of alternative productivity trends and harvest scenarios to inform collaborative management tables. We found declining abundances for 51% of coho populations since 2017, driven by productivity regime shifts associated with recent marine heatwaves. Although long-term coho recovery depended on future productivity trends, reduced harvest rates across U.S. and Canadian fisheries can improve short-term recovery prospects. While rebuilding coho will depend largely on whether productivity improves, harvest management remains one of the few tools available to provide a safe operating space for coho populations, and their fisheries, to adapt to ongoing ecosystem changes.

## Introduction

Marine species face increasingly unprecedented conditions from climate and ecosystem changes (Poloczanska et al. 2016; Bindoff et al. 2019). Migratory and anadromous species, like Pacific salmon, are particularly vulnerable to these changes as their life histories evolved to track the seasonal and geographic variation in climate and ecological regimes evident in both marine and freshwater environments (Arthington et al. 2016; Hardesty-Moore et al. 2018; Putman 2018). Climate-driven regime changes in productivity of exploited marine fishes can undermine conservation and erode the sustainability of critical social-ecological systems including fisheries (Vert-Pre et al. 2013; Poloczanska et al. 2016).

Over the past century, Pacific salmon have been managed into multiple conservation concerns – including an ongoing and widespread decline (Irvine and Fukuwaka 2011; Walters et al. 2019; Price et al. 2019). The reasons underlying these declines vary but common themes include multiple stressors from extensive climate, ocean, and freshwater changes integrated across a complex migratory life cycle (Lin et al. 2017; Cline et al. 2019; Wilson et al. 2022), degradation and loss of freshwater habitat (Gregory and Bisson 1997; McClure et al. 2008), and a legacy of overfishing (Achord et al. 2003). In 2014, for example, the North Pacific marine heatwave emerged from a suite of novel climate conditions (Di Lorenzo and Mantua 2016) that led to persistent changes in temperature, primary production, and trophic structure across the North Pacific (Suryan et al. 2021) and lowered marine survival for salmon populations from California to Alaska (Grant et al. 2019; Priest et al. 2021; PSC NBTC 2022). Ecosystem changes continue to impact salmon across the North Pacific and are associated with highly variable and non-stationary population dynamics that place tremendous pressure on their conservation and management (Peterman and Dorner 2012; Dorner et al. 2018).

In response to domestic fishery collapses, the Canadian Government has attempted to establish a foundation for precautionary management of wild Pacific salmon fisheries by enacting policies under the *Sustainable Fisheries Framework* (*SFF*) including the *Wild Salmon Policy* (*WSP*) and the *Fish Stocks Provisions* of the revised federal *Fisheries Act* (Holt et al. 2009; Price et al. 2017; Imhof et al. 2021). The guiding principles underlying the *SFF* are germane to fisheries management globally and include: (1) establish biological or ecosystem reference points that set low- and high-risk thresholds to be avoided (known in the *SFF* as the Upper Stock Reference [USR] and the Limit Reference Point [LRP], respectively); (2) conduct stock assessment that quantifies population status relative to those reference points; and (3) identify harvest strategy that allows some harvest while maintaining fish populations above the reference points with varying tolerance (i.e., high probability to avoid the LRP). Reference points used in stock assessments are typically based on adult abundances or biomass at maximum sustainable yield (*S*_*MSY*_) since population sizes below MSY are more likely to produce lower recruitment and lost fishing opportunities. In practice, British Columbia salmon management has routinely used 80% of *S*_*MSY*_ to define the USR before a population falls into a ‘cautious zone’, thereby warranting precautionary management actions that promotes recovery (we use the term USR interchangeably with 0.8*S*_*MSY*_ throughout). Similarly, practitioners have typically defined the LRP, before a population falls into a ‘critical zone’, as the spawner abundances that recover a population to *S*_*MSY*_ within a single generation (termed *S*_*GEN*_). However, neither reference point has been formally enshrined into policy nor mandated in practice. When applied at a Conservation Unit scale under the Wild Salmon Policy these two reference points delineate red, amber and green zones of biological “status” and are termed biological benchmarks (Holt and Irvine 2013).

Despite establishing a precautionary framework in Canada, efforts to conduct rigorous stock assessments and estimate reference points for wild Pacific salmon populations remain limited due, in large part, to the large number of salmon populations, scale mismatches between local population dynamics and larger-scale management units (e.g., Stock Management Units or Conservation Units; Wilson et al. 2023), and erosion of monitoring programs and associated budget cuts over time (Price et al. 2008, 2017; Walsh et al. 2020). This lack of monitoring data precludes quantifying reference points for many wild salmon populations, which can undermine precautionary fisheries management and the constitutionally protected rights of Indigenous communities for traditional food, social, and ceremonial (FSC) harvest. These challenges are well illustrated by coho salmon *Oncorhynchus kisutch* along the North and Central Coast (NCC) of British Columbia.

Coho in the NCC support vital commercial, recreational, and FSC fisheries, including for the Central Coast First Nations (CCFN). Fishers harvest coho salmon from hundreds of mixed and comigrating populations in the coastal environments of British Columbia and Southeast Alaska (Beacham et al. 2020; Priest et al. 2021). Despite their importance, monitoring and managing coho salmon populations remains challenged by difficult and remote access, and by limited resources and capacity throughout the NCC for monitoring and assessment at ecologically relevant scales (Atlas et al. 2021; PSC NBTC 2022). Local insights from Indigenous knowledge holders, Nation-led monitoring programs, and limited quantitative assessments since 2017 suggests an ongoing decline in coho abundances that coincided with the emergence of marine heatwave conditions in recent years (Grant et al. 2019; PSC NBTC 2022).

Here, we developed a Bayesian time-series model to evaluate spatial and temporal patterns in coho salmon population dynamics across 52 populations and six regions spanning the NCC. Overall, our goals were to estimate fishery reference points, assess current conservation status of NCC coho populations, and provide explicit management and ecological evaluations that illuminate a path forward for precautionary fisheries management. As several of us work directly for and with CCFN, our findings help to advance Indigenous-led fisheries management efforts and generates evidence-based advice for ongoing collaborative management agreements and conservation plans.

## Methods

### Spawner escapement and harvest data

We compiled time-series of adult spawner abundances and harvest rates from 52 coho salmon populations across the NCC (Figure 1) spanning a 41-year period from 1980 to 2020 and 16 Conservation Units (CUs), populations grouped on the basis of their geographic locations, genetics, life histories, and habitat conditions under the Wild Salmon Policy (Holtby and Ciruna 2007). While coho escapement data along the NCC goes back into the 1950s, we focused on 1980 and onwards owing to consistent survey methodologies and improved documentation (**Appendix S1:** Figure S1). The methods used to collect spawner abundance data varied across populations, and included weirs, overflight counts, stream walks, and mark recapture studies. We chose to focus on populations with relatively continuous monitoring over time across the period of interest. Harvest rates for coho salmon are highly uncertain and remain unmonitored across most of the NCC, except for three coded-wire tag (CWT) indicator populations on the Nass (Zolzap creek), Skeena (Toboggan creek), and Haida Gwaii (Deena creek). We used these indicator populations to develop estimates of total harvest rates from all sectors (Alaskan commercial and domestic Canadian commercial, recreational, and FSC) for coho across the NCC (Figure 2) based on methods outlined in English *et al*. (2018), with modifications detailed in

**Figure 1.**
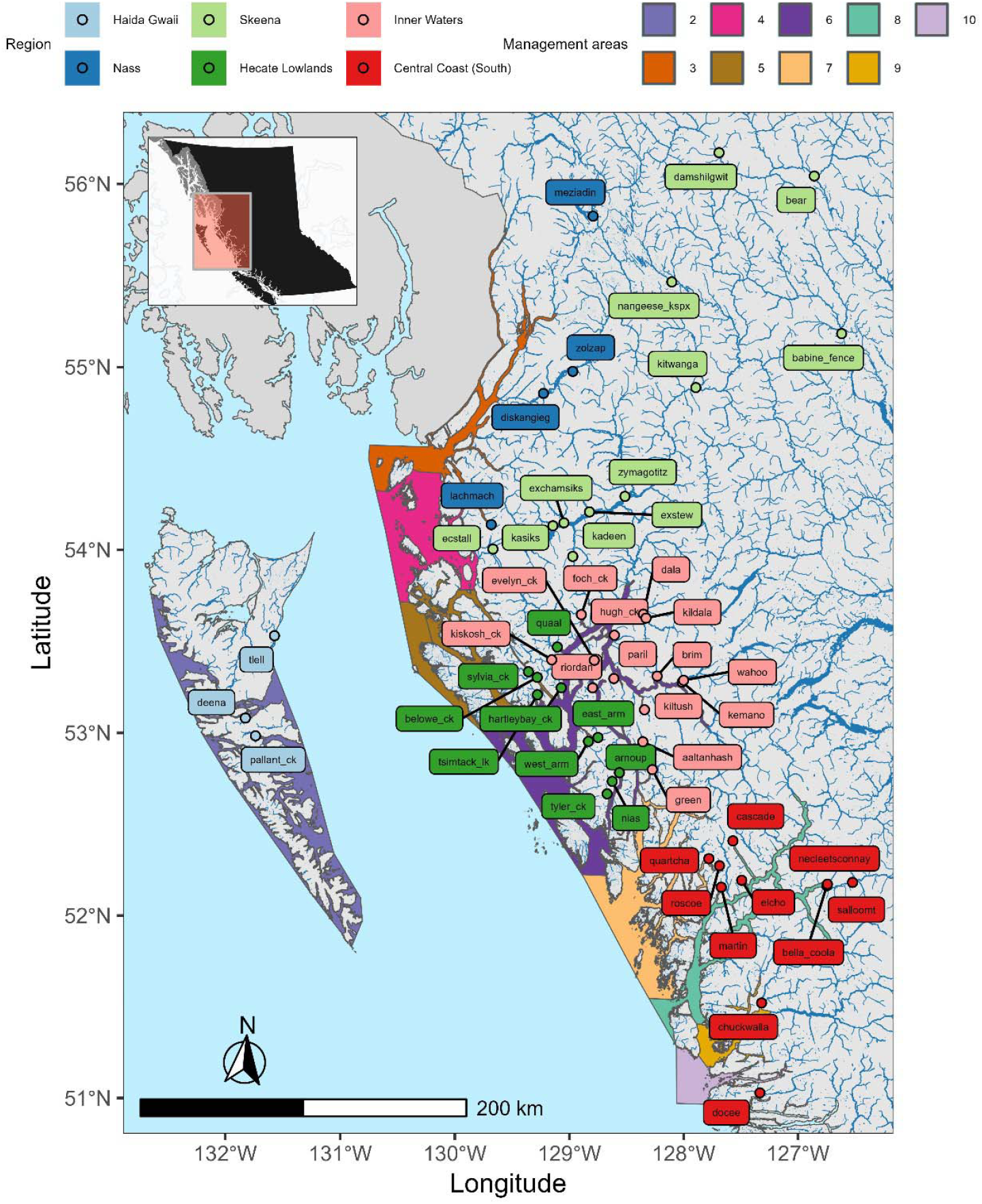
The Pacific Fisheries Management Areas and regional groupings for 52 coho salmon populations along the North and Central Coast of British Columbia.

**Figure 2.**
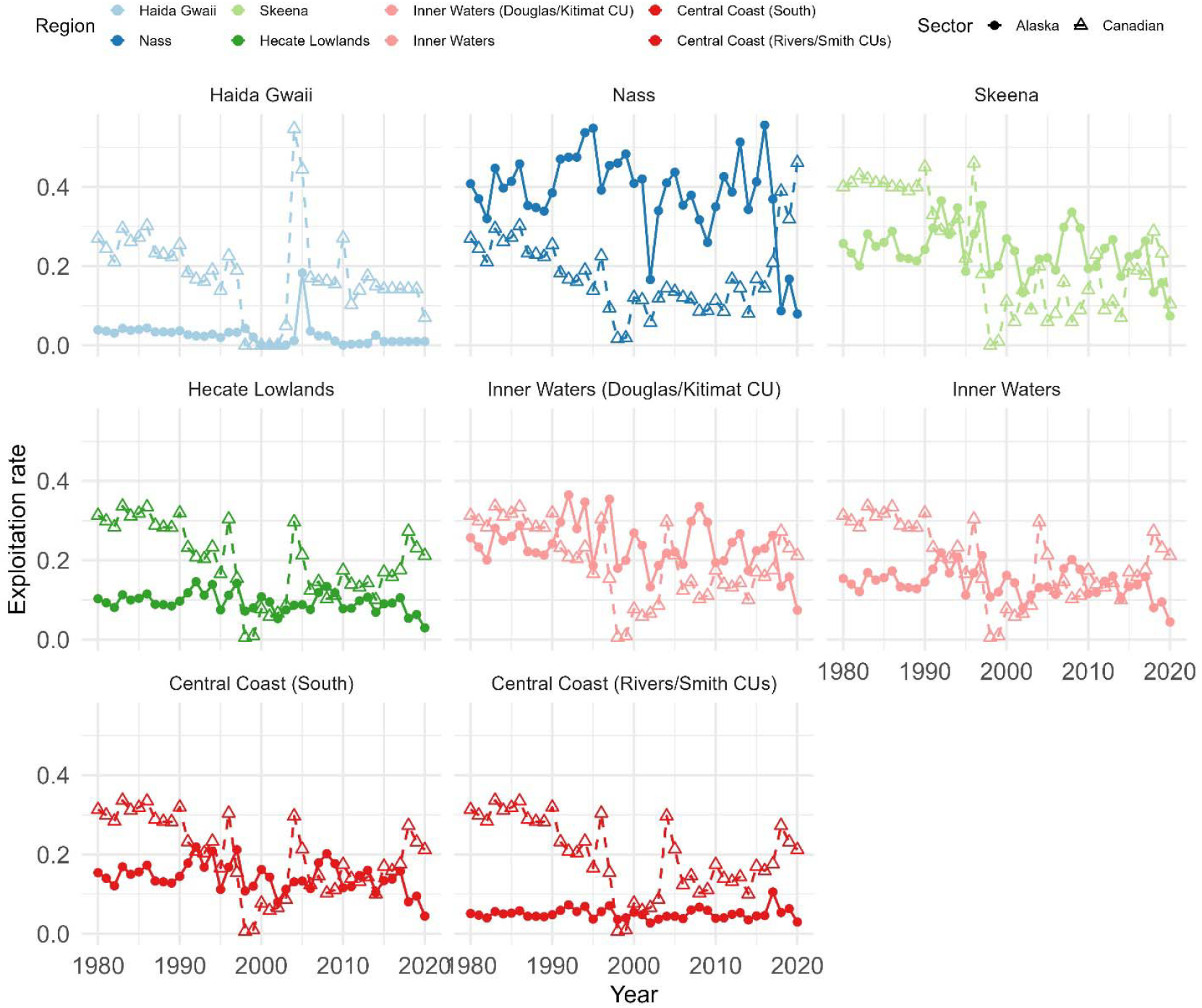
Trends in Canadian (open triangles) and Alaskan (filled circles) fisheries exploitation rates (ERs) for eight regions. Exploitation rates were estimated based on methods in English et al. (2018). Exploitation rates for coho populations in the Haida Gwaii, Nass, and Skeena regions were inferred from coded-wire tag recovery data from the Deena, Zolzap, and Toboggan Creek monitoring programs, respectively. Harvest rates for non-CWT regions were estimated as described in **Supplemental S1** and reflect the assumption that all regions are subject to the same average Canadian domestic exploitation rates.

## Appendix S1

### Coho population dynamics

The life cycle for each population was defined as a cohort of spawning coho adults and the number of subsequent adult recruits produced by that cohort. For our purposes, adult recruits were the sum of returning adults harvested in fisheries and those that successfully returned to their natal rivers to spawn (termed spawner escapement). Harvest was calculated based on harvest rates as modified from indicator populations (Figure 2) and spawner escapement, and returning fish were assigned to their parental brood year using age composition data in English *et al*. (2018). Coho salmon exhibit relatively simple age-structure compared to other salmonids. Accordingly, when populations lacked historical age composition data, we used the average proportions-at-age among populations (**Appendix S1:** Table S1), which assumed that freshwater rearing lasted 1-3 years (1.15 years on average) and a fixed 2-year period at sea prior to return as current spawning escapement counts include only adult spawners and not jacks (male coho that return after a single summer at sea).

#### Bayesian hierarchical model

We developed a hierarchical Bayesian multivariate state space model to describe non-stationary changes to the intrinsic productivity of 52 coho salmon populations within six regions among the NCC. Multivariate state space models are a useful way to analyse time-series data because they can simultaneously estimate shifts intrinsic productivity while accounting for the temporal dependency between observations (i.e., the autoregressive property), correlations between two or more time series (i.e., the multivariate property), and two sources of uncertainty: observation error and natural biological variation (i.e., the observation v. process states). Recruitment dynamics were analysed at the level of individual populations, rather than aggregating population dynamics across larger groups, because coho typically have low dispersal leading to density dependence regulating individual spawning populations rather than regional aggregates (Westley et al. 2013; Anderson et al. 2015). We modelled intrinsic productivity arising from a hyperprior that nested populations within six regional groups to account for common temporal trends among regions.

We assessed potential trends in recruitment productivity for coho salmon using a Ricker model (Ricker 1975; Brännström and Sumpter 2005) that describes a stationary density-dependent relationship between adult spawners *S* and subsequent recruitment *R*. The basic Ricker stock–recruitment model for population *i* follows:

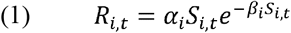

where *α*_*i*_ was the recruits produced per unit spawner at low densities (i.e., maximum intrinsic productivity) and *β*_*i*_ reflects the strength of density dependence such that 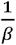 is the adult abundances that produce the maximum number of recruits. We linearized the Ricker model into a simpler regression-like form such that:

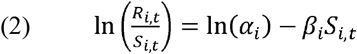

where 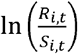 measures the observed recruitment productivity for population *i* in year *t*. Hence, a linearized Ricker model allowed us to quantify how intrinsic productivity (*α*_*i*_) and density-dependence (*α*_*i*_) jointly shaped recruitment productivity in NCC coho salmon populations.

The above eq. 2 assumes that the stock–recruitment relationship, defined by *α*_*i*_ or *α*_*i*_, remains constant through time, resulting in a stationary regime where recruitment dynamics respond solely to changes in adult densities or stochastic processes. We added a time-varying component (i.e., non-stationary) to eq. 2 to allow us to estimate persistent trends in coho productivity regimes, modelled as ln(*α*_*i*,*t*_), that varies for each population *I* at time *t* such that:

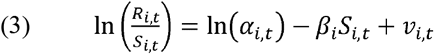

where *v*_*i*,*t*_ represented normally distributed errors with mean zero and standard deviation *σ*. Modelling recruitment as a time-varying process allowed us to estimate and detect trends in productivity regimes (Dorner et al. 2008; Wilson et al. 2022). Similar to Dorner *et al*. (2008), we evaluated for non-stationary changes in ln (*α*_t_) rather than shifting *β_t_* (or both). These types of time-series models are commonly used in stock assessments when productivity is believed to be temporally variable (Fisheries and Oceans Canada 2020a).

Annual variation in intrinsic productivity ln(*α*_*i,t*_) for each population *i* were then allowed to (co)vary among the six regional groupings *g* following a multivariate distribution such that:

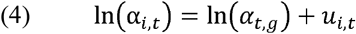

Where *u*_*i*_,_*t*_ represented normally distributed errors with mean zero and standard deviation *σ*_*α,g*_ (i.e., shared variance for populations within group *g*). If there was no evidence of time-varying productivity, then estimates for *u*_*i,t*_ in the process state would drop towards zero leading to a constant and stationary regime that only varies between groups. Specifically, we characterized changes in intrinsic productivity for each of the six regional groups using a multivariate normal distribution with a mean vector ln (*α*_t,g_) describing the expected intrinsic productivity for populations within six regional groupings at time *t* and a variance-covariance matrix Σ_p_ modelling process error within and between these groups. Σ_*p*_ The hierarchical regional averages ln (*α*_t,g_) followed an autoregressive time-varying process state such that:

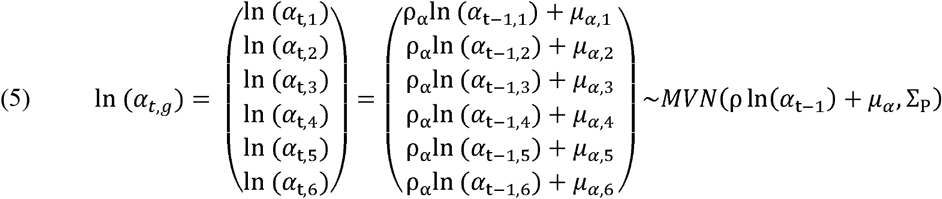

where ρ was the temporal autocorrelations between successive time-steps and a long-term mean *μ*. The variance-covariance matrix (Σ_P_, a 6x6 matrix) was drawn from a vague inverse-Wishart prior. Spawner abundances followed similar temporal dependencies between time-steps (but no regional hyperpriors) such that:

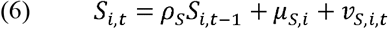

where *v*_*s,i,t*_ represented lognormal errors, ρ was temporal autocorrelation, and mean *μ*_*s*_. Model shape parameters, such as annual productivity *α* or trend terms, were estimated by fitting equations 3-6 to observed recruitment productivity (eq. 3). The dynamics represented in eq. 5-6 presumed that intrinsic productivity (or spawner escapement) would be expected to correlate to the previous year’s productivity (or escapement) and will likely return to a long-term mean *μ*_*α*_ (or *μ*_*s*_). Below, we relaxed and tested the assumption of productivity or spawner abundances reverting to a long-term mean.

#### Reference points

Current fishery policies in Canada establish a need for setting biological reference points to determine population status and assess conservation risk. Two biological reference points are typically used in practice at the scale of the Conservation Unit: (1) 0.8*S*_*MSY*_, which corresponds to the Wild Salmon Policy upper reference point (Holt et al. 2009; Holt and Bradford 2011) and which for the purposes of this paper we considered synonymous with the USR (delineating ‘healthy’ from ‘cautious’ status) and (2) *S*_*GEN*_, which corresponds to the Wild Salmon Policy lower reference point and which we considered synonymous with an LRP (delineating ‘cautious’ from ‘critical’ status). These reference points revolve around the concept of a stationary (i.e., non-trending) production regime that can be used to describe a population’s maximum sustainable yield, typically estimated using numerical methods from the underlying production model. We followed Scheuerell (2016) to estimate spawner abundances at maximum sustainable yield *S*_*MSY*_ for population *i* such that:

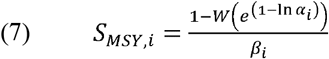

where W was the solution to Lambert’s function and *α_i_* and *α_i_* were the long-term average demographic parameters estimated from our Bayesian hierarchical time-series model fitted to 1980-2020 data to approximate a presumed stationarity regime. We then used *S*_*MSY*_ to estimate *U*_*MSY*_ (the harvest rate at MSY) and *S*_*GEN*_ using numerical methods following Holt *et al*. (2009). We added a third reference point unrelated to MSY based on a population’s average spawner abundances between 2000-2015 (termed *S*_*Baseline*_), as this time-period corresponded to a We added a third reference point unrelated to MSY based on a population’s average spawner widespread recovery and improved harvest from the Central Coast First Nations following a three-year reduction of domestic coho harvest rates from 1998-2000. We derived posterior distributions for each of 0.8*S*_*MSY*_ and *S*_*GEN*_ for each coho population using posterior distributions of *α* and β values from preliminary model fits to the spawner-recruitment datasets. For simplicity, our analyses only used the posterior mean for each reference point. We also demonstrate how reference points fluctuate temporally as the underlying demographic parameters, like *α*, change through time.

#### Forward simulations and fishery risk analysis

We fitted eq. 3-6 to empirical observations of 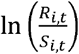 and then forward simulated recruitment dynamics five generations (20 years) into the future under five alternative harvest scenarios and three assumptions about future productivity trends. For our harvest scenarios, future recruitment from a spawning cohort at time *t* followed:

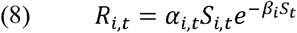

where the abundance of returning adult spawners *A*_*i,t*_ depended on the proportion-at-age in the ocean *p*_*i,a*_ of that population (from 3-5 years):

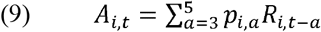

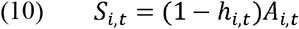

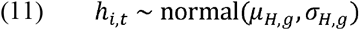

where *h*_*i,t*_ was the harvest rate (truncated positive) experienced by population *i* at time *t, *μ**_*H,g*_ was the scenario-specific mean annual harvest rate and *σ*_*H,g*_ was the standard deviation in annual harvest rates for populations within region *g* from the past ten years (2009-2019). We did not apply the log-normal bias correction to eq. 8, which allowed for the risk assessments in forward simulated dynamics to reflect the posterior median recruitment rather than the posterior mean. We evaluated five harvest scenarios: (1) no harvest, (2) harvest continues at the recent ten-year average, (3) 50% reduction of BC harvest, (4) 50% reduction of Alaska harvest, and (5) 50% reduction of both BC and Alaska harvest. These scenarios capture the full range of potential management interventions from status quo to domestic reductions (in BC or Alaska) to internationally coordinated reductions (in BC and Alaska) to complete cessation of commercial, recreational, and FSC fisheries.

Our forward projections then evaluated outcomes across three alternative future states of nature as indexed by alternative assumptions for non-stationary shifts in the maximum intrinsic productivity *α*_*t,g*_ (**Appendix S1:** Figure S2). The first scenario simulated maximum intrinsic productivity returning to a long-term mean such that

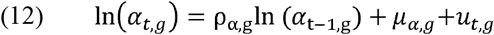

where ρ_α,g_ was the temporal correlation between productivity at time *t* and time *t-1*, *μ*_*α,g*_ represented average productivity for group *g*, and 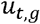 were process errors estimated from a variance-covariance matrix (eq. 6). The AR(1) process implied by eq. 12 reverts to a long-term mean of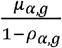. The second scenario simulated maximum intrinsic productivity and adult spawners continuing along a random walk such that:

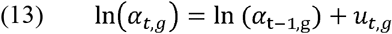

The third scenario simulated maximum intrinsic productivity following long-term linear trends such that:

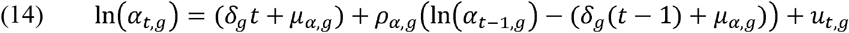

where *δ*_*g*_ was the annual rate of change in average productivity *μ*_*α,g*_ for group *g* and *ρ*_*α,g*_ was the temporal correlation between time-steps. Note that eq. 14 models a linear trend in the long-term average productivity where the expected productivity was centered on the long-term trend plus a proportional deviation from the previous autocorrelated time-step to account for the temporal dependency among consecutive years (**Appendix S1:** Figure S2).

We used forward simulations of alternative scenarios to quantify risks for each population, each regional group, and the NCC aggregate, with risks measured in two ways. First, we quantified the frequency of the posterior samples that populations fell below each of the above fishery reference points (0.8*S*_*MSY*_, *S*_*GEN*_, and *S*_*Baseline*_). Second, we quantified population status measured as the ratio between simulated abundances and these reference points across all projection years.

#### Inference & model diagnostics

We estimated the Bayesian model on a joint posterior using four Markov Chain Monte Carlo (MCMC) chains in JAGS version 4.3.1 run via R version 4.3.1 with *rjags* and *run*.*jags* packages (Plummer 2003; Denwood 2016; R Core Team 2023). Each chain took 1,500 posterior samples for a total of 6,000 samples, and we thinned every 200 samples after a warmup period of 125,000 samples. We used several complementary methods to diagnose model suitability. MCMC chain convergence was inspected visually on trace plots. In addition, we ensured effective sample sizes were >1,000 for each parameter (Gelman et al. 2013). We used the Gelman-Rubin diagnostic test on each parameter to determine whether independent chains converged to a common posterior mode, with potential scale reduction factors (PSRF) < 1.1 suggesting convergence. We then used graphical posterior predictive checks to test for model misspecification by comparing the predictive distribution of adult escapement (simulated from the posterior sample for each observation) to observed population dynamics. Last, we inferred the weight of evidence for a given assumption on forecasted productivity by calculating the absolute median error between predicted and observed escapement for 2017-2020 return years for the three productivity models. We used absolute median errors because it is robust to outliers that can emerge from forecasting population dynamics under lognormal errors.

#### Marine climate – productivity associations

We used an index of marine heatwaves for each of the six regional groupings to explore the potential role of climate change on coho salmon. First, we derived ocean entry locations for each region by averaging the ocean entry coordinates for coho populations within each region. We then compiled gridded sea surface temperature (SST) data from NOAA ERSST (Boyin et al. 2017; Malick 2019) to calculate annual SST anomalies within 200km of each region’s ocean entry point. We then averaged SST anomalies at both an annual (January–December) and summer (May–September) time-period for each region to reflect the marine climate conditions that coho would experience during their first year at sea. Last, we used linear regression to relate annual SST anomalies to the posterior mean estimates of the regional time-varying productivity two years earlier (presuming an average of one-year spent in freshwater) to explore the associations between marine heatwaves and coho population dynamics (Mueter et al. 2002).

### Results

#### Trends in coho populations

In general, we found that NCC coho populations have undergone widespread declines in recent years (**Appendix S1:** Figure S1). Since 2017, spawner abundances declined by 39% on average compared to their contemporary baseline (*S*_*Baseline*_). Local monitoring indicated that 39 of 52 populations were below *S*_*Baseline*_ from 2017 to 2020 while 4 populations remained above; 9 populations were not monitored in recent years. The magnitude of population changes varied among regional groups. For example, Hecate Lowlands populations declined the most (mean: - 47%, range: -88–66%) while Nass River populations declined the least (mean: -13%, range: -40– 17%).

Intrinsic productivity of NCC coho populations exhibited large fluctuations with several boom-and-bust cycles since 1980 (Figure 3). For example, a period of increased productivity across several regions (e.g., Inner Waters in Figure 3), was associated with a rapid recovery due to high recruitment years, which coincided with reduced harvest in the fishery after the 1997 ‘coho collapse’. Since 2015, however, intrinsic productivity (ln *α*) for all six regions has substantially declined relative to their contemporary baselines with productivity nearing or exceeding record lows for three regions: Hecate Lowlands (posterior median percent changes of -74%; 95% credible intervals [CI]: -87 – -54%), Central Coast – South (-44%; CI: -68 – -2%), and Skeena Watershed (-69%; CI: -78 – -56%). Populations among the Central Coast Inner Waters changed by -45% (CI: -55 – -34%) – a large decline, but not the record low that occurred in the early 1990s. Estimated productivity trends were more moderate for coho populations among the Nass Watershed (-26%; CI: -45–4%) and were highly uncertain for populations among Haida Gwaii (-12%, CI: -74–114%).

**Figure 3.**
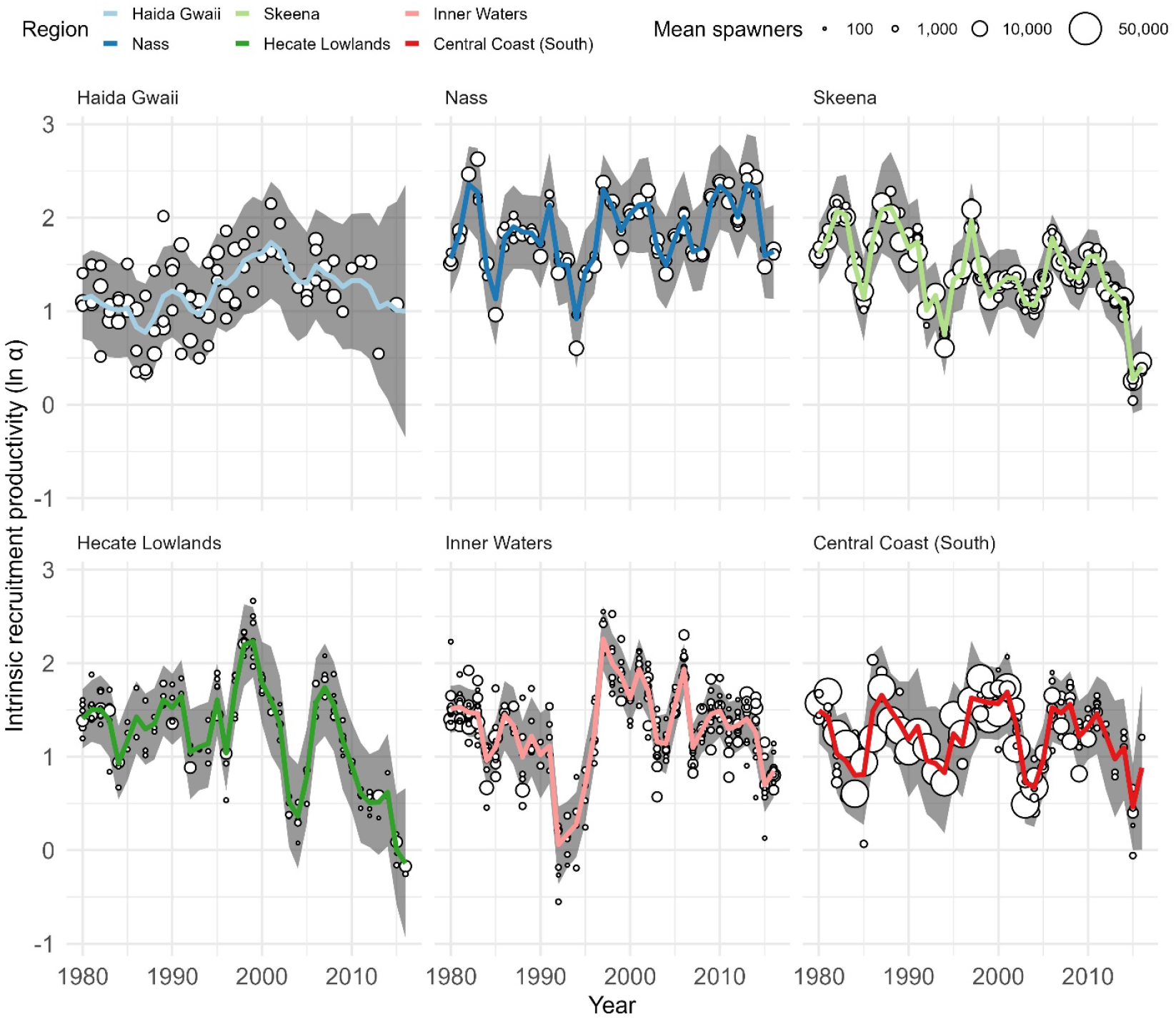
Trends in the intrinsic productivity (ln *α*) from 52 coho salmon populations (points indicate population-specific posterior median estimates) within six regions of the NCC (lines indicate region-specific posterior median, and the shaded regions indicating 95% credible intervals). The size of each point indicates the long-term average abundance for each population.

#### Current Population Status

Many coho populations were below conservation targets in recent years (Figure 4), while seven populations lacked monitoring data necessary to determine status since 2017. Of 45 monitored populations, 11 fell below their upper stock reference point (0.8*S*_*MSY*_) in each year they were monitored, and 34 populations were below in at least one year between 2017 and 2020 (Figure 4) suggesting ‘cautious’ status. Ten coho populations fell below *S_GEN_* in at least one year and two populations fell below in each year, placing them in or nearing ‘critical’ status. Not all populations declined in spawner abundances. For example, while 10 populations were below both 0.8*S*_*MSY*_ and *S_Baseline_*, 11 populations were above both reference points (Figure 4) and may be considered ‘healthy’ under current policy.

**Figure 4.**
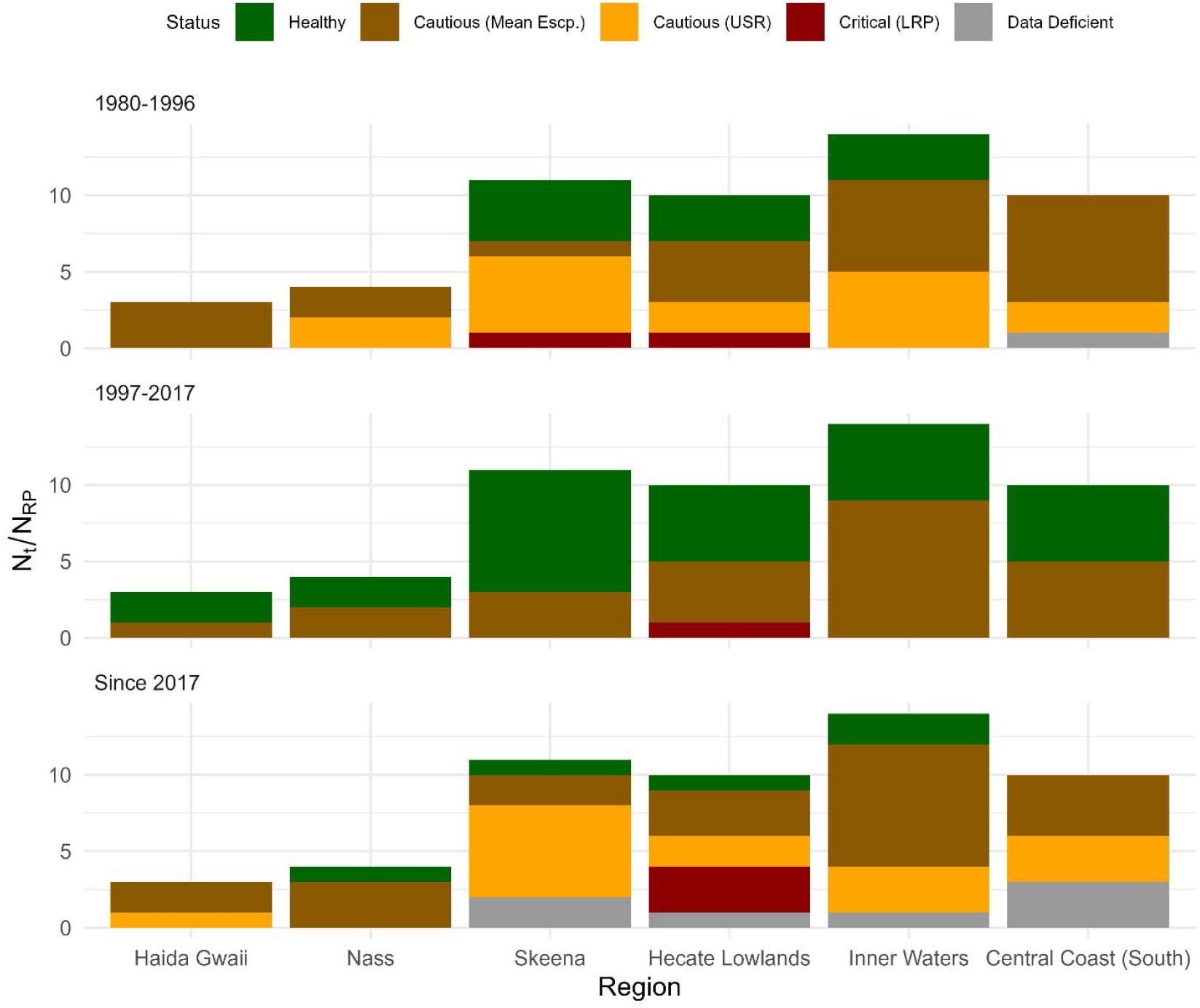
Population status of coho salmon in the North and Central Coast of British Columbia among three different time-periods. Population status indicated by posterior mean estimates of spawner abundance relative to estimated reference points. Healthy status is indicated if spawner abundance was above both the Upper Stock Reference Point and the Contemporary Baseline. Cautious status is indicated if spawner abundance was below either the Upper Stock Reference Point or the Contemporary Baseline. Critical status is indicated if spawner abundance was below Limit Reference Point. Data Deficiency is if there was no observed spawner abundance for the population during that time-period.

The role of fishing in driving declines in coho populations status varied over the past 40 years (**Appendix S1:** Figure S3). In the 1990s, many populations were overfished and undergoing overfishing 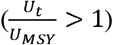, but fisheries management generally reduced harvest beginning in 1997 (**Appendix S1:** Figure S3) leading to mixed conservation outcomes (Figure 4). For instance, coho populations in the Skeena or the Central Coast (South) regional groupings saw an increased frequency of populations in Healthy status zone after harvest reductions in 1997, while the Hecate Lowland or Inner Waters region continued to see a relatively large proportion of populations remain in Cautious or Critical status zones (Figure 4). Time-varying productivity drove non-stationary changes in reference points, like *U*_*MSY*_, likely underestimating conservation risks in regions with large productivity declines (**Appendix S1:** Figure S3).

#### Marine climate and productivity

Based on post-hoc regression analyses, we found regional differences in the response of intrinsic productivity to warmer ocean climates indicated by annual and summer SST anomalies (Table 1; **Appendix S1**: Figures S8 and S9). For example, the posterior mean estimates of intrinsic productivity for the Nass and Haida Gwaii regions had weak or uncertain associations with either SST metric (Table 1). However, SST anomalies were negatively associated with intrinsic productivity for more southern regions of coho salmon including populations in the Skeena, Central Coast South (albeit uncertain effect), Hecate Lowlands, and Inner Waters regional groups.

**Table 1.**
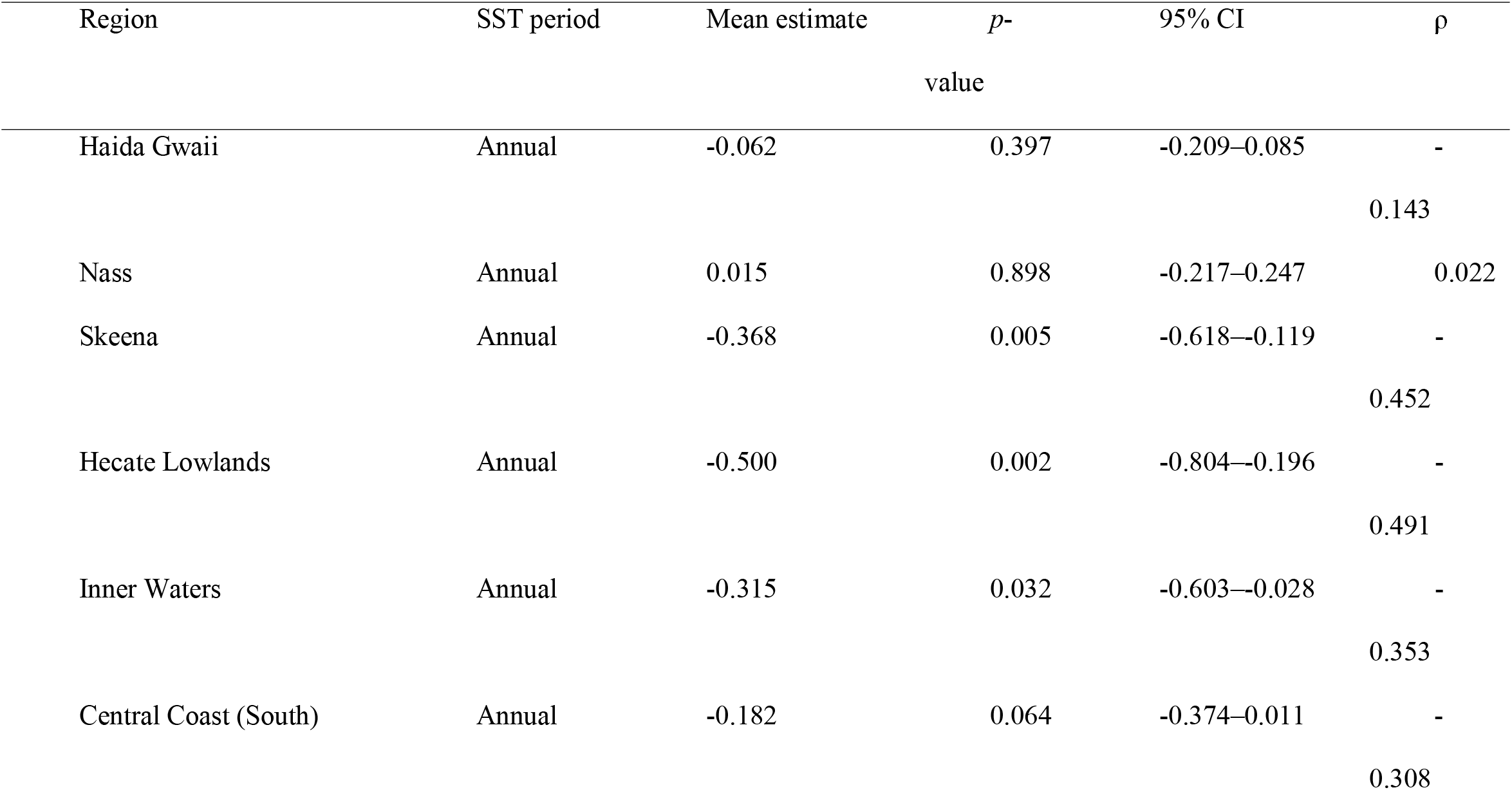
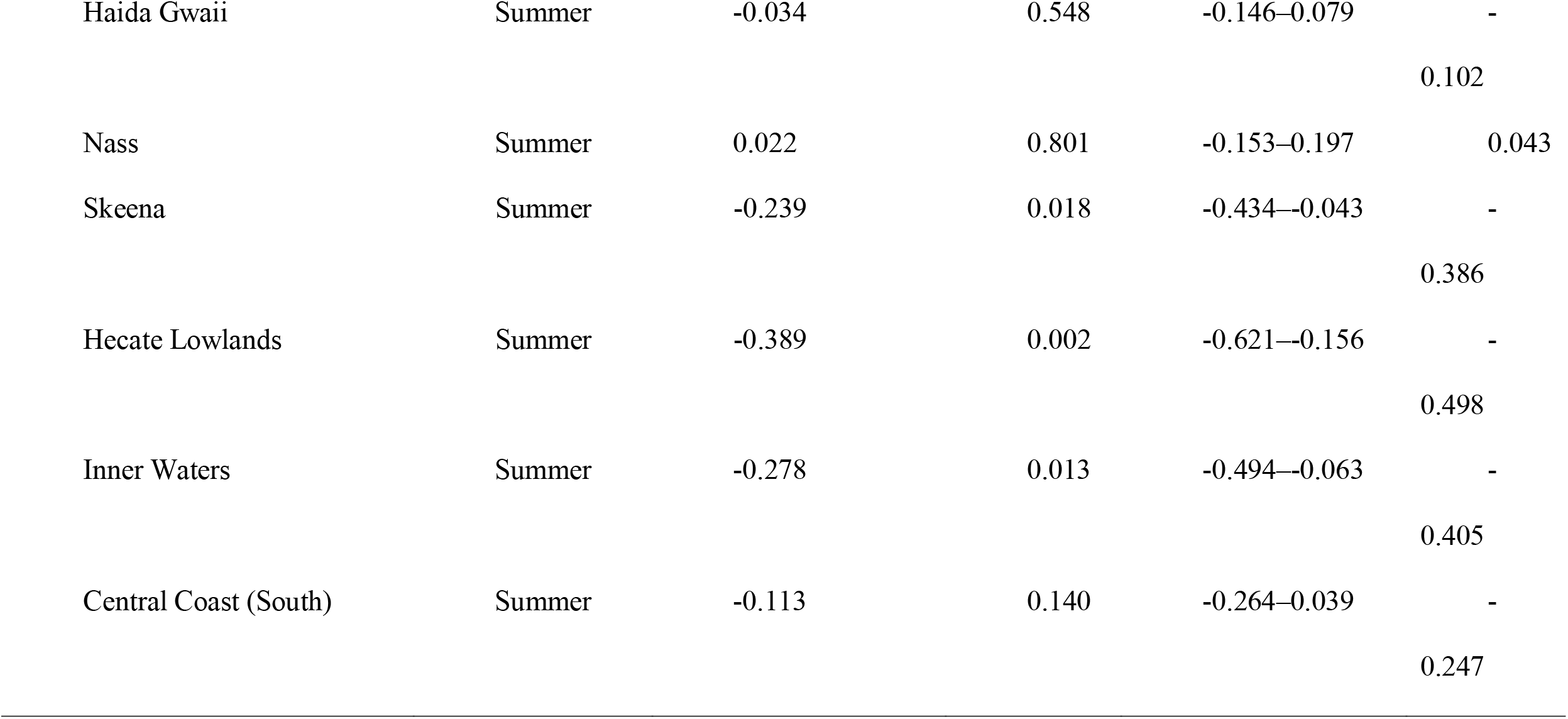
The mean estimated effect, *p*-value, 95% Confidence Interval [CI], and correlation (ρ) between sea surface temperature anomalies (annual and summer) and posterior mean estimates of intrinsic productivity ln(*α*) for six regional groupings of coho salmon populations based on regression analyses.

#### Forward simulations and recoveries

Forward simulations indicated that coho population recoveries depended upon future productivity trends and harvest management (**Appendix S1:** Figure S4; Appendices S2–S4). In aggregate, models with declining productivity trends improved the predictive accuracy of 2017-2020 spawner abundances by 3.6% compared to models with productivity returning to a long-term mean, but forecasted trends varied among populations and regions (Figures 5 and S5). Across the entire NCC, fishery risks under status quo management varied among the productivity models (Figures 5 and S6). Overall, the ‘mean reverting’ model, where intrinsic productivity returned to contemporary baselines, led to improved abundance and returned many populations into ‘healthy’ status (posterior frequency of future population-years: 7%≤LRP, 29%≤USR, and 65%≤baseline abundances). However, both the ‘random walk’ and ‘trending’ productivity models generally predicted declining productivity and led to lower abundances and increased conservation risks (‘random walk’: 38%≤LRP, 59%≤USR, and 78%≤baseline abundances; ‘trending’: 44%≤LRP, 62%≤USR, and 78%≤baseline abundances). The extent of these risks varied within and among regions (Figures 5 and S7). Across productivity scenarios, populations among Haida Gwaii, Nass Watershed, and the Central Coast – Inner Waters would likely maintain above their respective USR but the risks that populations fell below their LRP increased for ‘random walk’ and ‘trending’ scenarios (Figures 6 and S7). Conversely, large risks remained for populations among Hecate Lowlands and Central Coast South even under the ‘mean reverting’ model (Figures 6 and S7).

**Figure 5.**
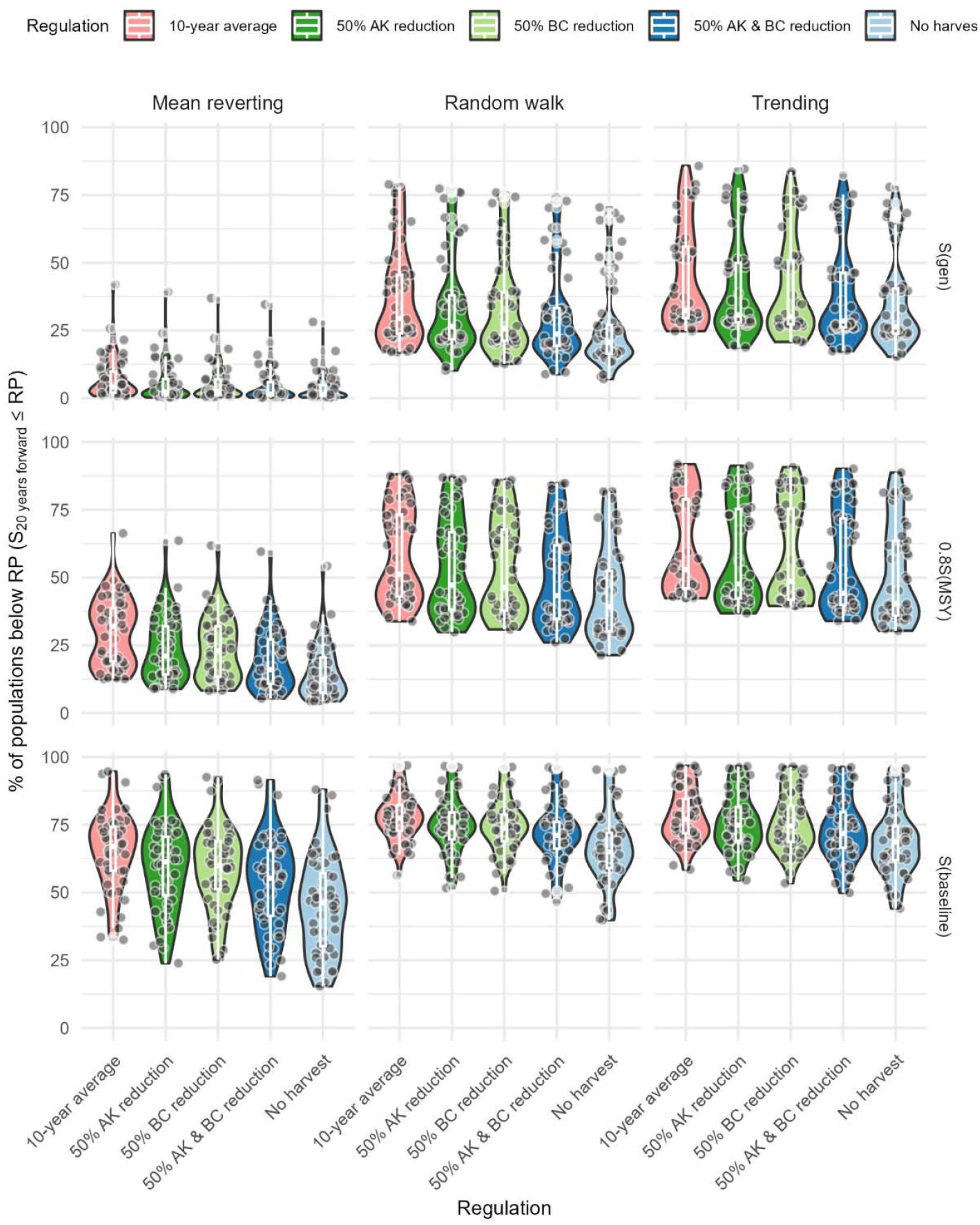
Posterior frequency distributions that future years of NCC coho salmon populations (points) fell below either of three fishery reference points under forward simulations of three productivity and five harvest management scenarios at the aggregate scale. Each point represents an individual population with that region.

**Figure 6.**
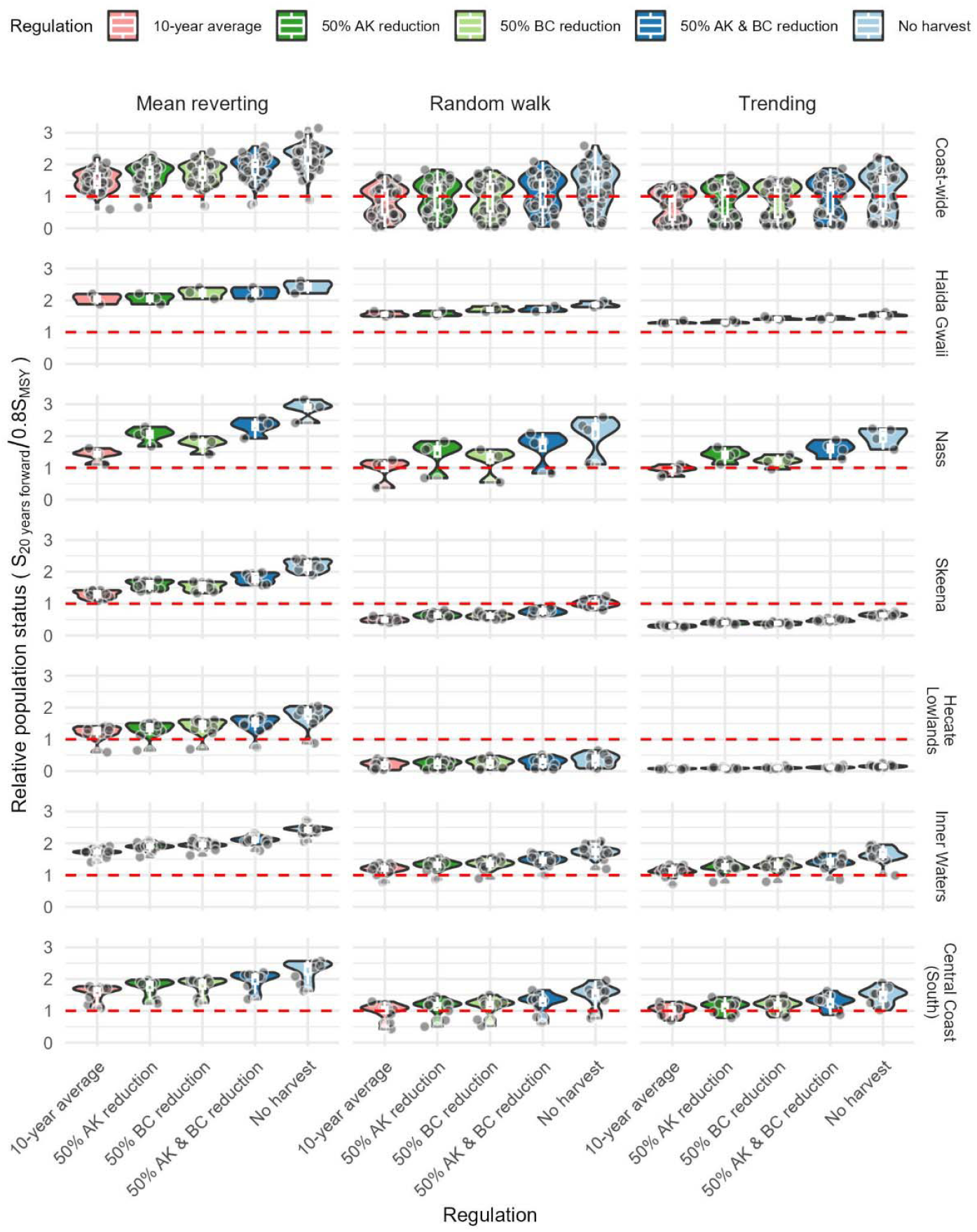
Posterior distributions of the relative population status 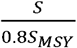 of NCC coho salmon populations under forward simulations of three productivity and five harvest management scenarios at aggregate (top panel) and regional scales. Each point represents an individual population with that region.

The forward simulations suggest precautionary fisheries management (i.e., reductions in mixed-stock harvest) would benefit future coho recoveries, but the magnitude of conservation benefits are expected to vary among productivity scenarios, regions, and risk metrics. For example, a 50% reduction in exploitation rates from all fisheries (Canadian and Alaskan) was projected to reduce the frequency of future population-years falling below their USR by 10% (≤*S*_*Baseline*_: 12%) under the ‘mean reverting’ productivity model, 8% (≤*S*_*Baseline*_: 7%) under the ‘random walk’, and 6% (≤*S*_*Baseline*_: 4%) under the ‘trending’ model. In contrast, closing all fisheries was projected to reduce the frequency of falling below their USR by 20% (≤*S*_*Baseline*_:20%), 12% (≤*S*_*Baseline*_:12%), and 10% (≤*S*_*Baseline*_:9%) for the ‘mean reverting’, ‘random walk’, and ‘trending’ models, respectively. The relative population status after 20-years of forward simulations varied among regions and depended on productivity trends and management scenarios (Figures 6 and S6). For example, reducing harvest by 50% among both fleets would improve population status by 28% under the ‘mean reverting’ productivity model, 32% under the ‘random walk’, and 31% under the ‘trending’ model. In general, regions with larger declines in productivity, like Hecate Lowlands, benefited more from precautionary management and their status shifted from near or below fishery reference points (under status quo harvest) to above fishery reference points under fishery reductions (Figure 6), but this effect varied among productivity scenarios.

## Discussion

Our findings provide new insights into recent declines of coho salmon along the North and Central Coast of British Columbia and their prospects for rebuilding. We found four main takeaways after evaluating 40-years of population dynamics and forward simulations of ecological and management scenarios. First, we found strong evidence that non-stationary productivity regimes contributed to large declines in coho salmon, altering their ability to support fisheries (Figure S3). Based on recent estimates of spawner abundances since 2017, we found that 36% of NCC coho populations were in a ‘cautious zone’ (i.e., above their LRP but below their USR) and 15% of populations were in a ‘critical zone’ (i.e., below their LRP) – which is consistent with a recent assessment of status for coho salmon Conservation Units across the NCC (Price et al. 2017). Second, we found that precautionary fisheries management that reduces harvest from both Canada and Alaska fisheries can benefit coho returns. Third, we found that the long-term abundance of coho largely depended on whether productivity improves in the future. Last, we found that different reference points convey different insights into population status. Reference points based on MSY, which assume stationary dynamics no longer apparent in an era of rapid climate change, may be overly optimistic compared to long-term ecological baselines such as those reflected by Indigenous knowledge (Jardine 2019; Lamb et al. 2023).

Pacific salmon face unprecedented environmental conditions driven by climate changes in the North Pacific. Overall, we found that coho spawner abundances declined by 51% across the North and Central Coast of British Columbia and was driven by coast-wide declines in intrinsic productivity, with record or near-record low productivity observed for several regions. Coho spawner abundance has high natural variability and is subject to cyclical dynamics driven by variation in climate (Bradford 1995; Logerwell et al. 2003; Briscoe et al. 2005). However, the timing of recent declines appears strongly associated with impacts by recent marine heatwaves that have persisted in varying degrees of intensity since 2014 (Di Lorenzo and Mantua 2016; Mantua 2019; Suryan et al. 2021) and continue to impact salmon ecosystems across British Columbia (Grant et al. 2019; PSC NBTC 2022). Unfortunately, this paints a more pessimistic outlook on the pathway for coho salmon recoveries along the NCC – a likely reason underlying our stronger empirical support for recent trends in intrinsic productivity.

Preliminary explorations of associations between nearshore ocean temperatures and regional trends in coho salmon productivity revealed apparent geographic differences in the role of warm ocean climate on annual estimates of regional coho productivity. Linear models relating SST and intrinsic productivity for Nass and Haida Gwaii populations had near zero slopes, suggesting limited effects of warm ocean on survival and recruitment. However, Skeena and three regional groupings of Central Coast coho populations (Inner Waters, Central Coast South, and Hecate Lowlands) all showed evidence of negative associations between warm nearshore ocean conditions and intrinsic productivity. There was also apparent divergence across these regions in productivity trends, with evidence for more stable productivity among Nass and Haida Gwaii populations during recent marine heatwaves. These findings are consistent with research that has demonstrated differential responses to climate changes among salmon populations with more northern and more southerly salmon populations of multiple species, possibly due to differing oceanographic processes driving productivity between the northern and southern bifurcations of the North Pacific Current (Mueter et al. 2002; Malick et al. 2017; Litzow et al. 2019).

The emergence of non-stationary shifts in intrinsic productivity challenges conservation as managers must navigate a future with no historical analog. Traditional salmon management by colonial governments has been informed by retrospective assessments of broad-scale abundance data aggregated among populations that establish equilibrium-based reference points (Holt et al. 2009; Price et al. 2017). However, ongoing climate and ecosystem shifts will drive non-stationary marine and freshwater productivity regimes, suggesting that time-varying reference points, like *U*_*MSY*_, may be more appropriate (**Appendix S1:** Figure S3). These non-stationary dynamics are a common feature among Pacific salmon (Szuwalski and Hollowed 2016; Malick et al. 2017; Litzow et al. 2018; Wilson et al. 2022) and force managers to aim at moving targets (Ingeman et al. 2019). In addition to environmental changes, regime changes can emerge from social or economic changes. For example, previous NCC coho declines in the 1990s were associated with large improvements to fishing technologies, including electronic navigation aids and GPS, that improved capture efficiencies and led to more widespread fishing activity throughout the NCC (Mike Reid, personal communication).

Unfortunately, such rapid productivity changes may go unnoticed for NCC coho because of declining monitoring efforts and a lack of established reference points, in particular at locally and regionally relevant scales (Price et al. 2017). Despite the socio-cultural and economic importance of coho salmon, estimating fishery reference points for precautionary management of NCC coho salmon remains challenged by limited spawner enumeration and patchy catch monitoring. These limitations require researchers and managers to rely on numerous assumptions related to expansions of visual count data, population age structure, and harvest rates across different regional population groups. While the underlying trends in abundance and productivity we document across regions are likely robust, estimating individual population status and managing fisheries will require improved catch and spawner escapement monitoring for culturally and ecologically significant populations.

Recent collaborative governance agreements between federal and First Nations fisheries managers demonstrate the need to improve monitoring that can navigate these concerns and develop precautionary and adaptive management (Atlas et al. 2021). In some cases, current policy and institutional barriers limit such opportunities. For example, under the Salmon Allocation Policy (currently undergoing review in 2023-25), federal managers are unable to reduce recreational harvest (a substantial portion of domestic harvest) without first identifying a conservation risk or closing commercial fisheries (DFO 1999). This action has not been undertaken in NCC coho fisheries since 1999. Yet, establishing reference points and taking a more active role in managing fisheries in response to annual variations in productivity will be critical if Pacific salmon are to continue supporting Indigenous, recreational, and commercial fisheries into an uncertain future (Whitney et al. 2020).

Reference points used to measure risk under current policy frameworks often lack historical baselines that reflect culturally meaningful abundances and may inadvertently promote risk rather than precaution (Frid and Atlas 2020; Lamb et al. 2023). Reference points such as *S*_*MSY*_ are designed to maximize fishery yields under stationary conditions, but populations can exhibit non-stationary dynamics that lead to time-varying reference points. These time-varying refence points may serve to reinforce or accelerate declining abundance in instances where productivity is low or undergoing a declining trend. Managers lean on limit reference points such as *S*_*GEN*_, to curtail harvest when populations fall below critical thresholds – in this case, *S*_*GEN*_ denotes the spawner abundances necessary to recover the population to *S*_*MSY*_ within one generation. By definition, a USR of 0.8*S*_*MSY*_ is 20% lower than *S*_*MSY*_– a common reference point used in other jurisdictions (e.g., the US Magnuson-Stevens Act; U.S. Department of Commerce 2007). Reference points based on MSY, including both USR and LRP, allow for fisheries to reduce populations relatively low abundances compared to historical baselines before management would reduce harvest (Frid et al. 2023). Indigenous communities, including the CCFN, would prefer more precautious reference points (e.g. ≥*S*_*MSY*_) be established that better incorporates Indigenous values and promotes more robust conservation, recovery and local abundance by placing greater restrictions on harvests, for a given spawner abundance, than other less biologically precautionary reference points like 0.8Smsy (Frid and Atlas 2020). One such example could be our third reference point, *S*_*Baseline*_, the mean spawner abundances from 2000– 2015, a recent time period when coho salmon provided reliable benefits to Indigenous, recreational, and commercial fisheries and when harvest rates were relatively consistent (PSC NBTC 2022). Similar reference points based on recent time-periods of suitable fishery catches have been used in the draft rebuilding plan management of Pacific herring in Haida Gwaii (Haida Nation et al. 2022).

Precautionary fisheries management can help coho salmon conservation but requires effectively implemented collaborative governance and coordination between multiple sectors and jurisdictions including Indigenous, Canadian, and US fishery managers. Harvest rates based on indicator populations suggests a historical legacy of overfishing for many populations and regions of the NCC prior to the 1990s, when the collapse of coho fisheries forced management to reduce harvest, restructure the commercial fleet, and implement precautionary management to protect less productive populations (Fisheries and Oceans Canada 1999, 2020b; PSC NBTC 2022). However, harvest remains unquantified for most coho salmon populations, and there is likely considerable variability in harvest rates that is uncaptured by the three CWT programs. While the three CWT programs suggest relatively modest harvest in recent years, how harvest from diverse commercial, recreational, and Indigenous fishers varies among coho populations depends on their marine distribution and patterns in migration timing (Beacham et al. 2020). For example, troll fisheries in Southeast Alaska may disproportionally harvest outer coastal populations of coho salmon due to their prolonged over-summer feeding and residency in coastal waters (Shaul et al. 2014). These insights depend upon reliable harvest estimates, which has been challenging for mixed stock fisheries like coho salmon in the NCC. Inaccurate harvest estimates from unmonitored populations can bias population productivity estimates and risk assessments, jeopardizing recovery prospects and the ability of salmon populations to sustain fishery opportunities. However, the productivity values we report here are well within the range of values observed for coho populations (Bradford et al. 2000), and group-level productivity trends appear reliable since they reflect aggregate dynamics that integrates variability within and among populations.

The contrasting recovery scenarios we highlight illustrate the sensitivity of future fishery opportunities to ongoing climate perturbations and changes in marine and freshwater habitat conditions. Although harvest may not be the main driver of declines among wild coho salmon populations, reducing harvest rates may be one of the only tools available for management to try to increase the safe operating space for coho into the future (Dearing et al. 2014; Carpenter et al. 2017). Coho salmon are capable of high productivity and rapid recovery during periods of favorable environmental conditions but increasing climate variability underscores the need for improved monitoring, particularly to provide high spatial- and temporal-resolution information on freshwater and marine survival, environmental indicators, fishery catches, and returning spawner abundances. Hence, harvest reductions may be a pragmatic choice to bolster the long-term chances of rebuilding coho populations. In an uncertain future, adaptive and precautionary management will be essential for balancing fishery opportunities while limiting conservation risks and fostering adaptation and resilience among wild salmon (Mantua and Francis 2004; Schindler and Hilborn 2015).

## Acknowledgements

We thank the many Charter Patrol and First Nations fisheries programs that contributed data to these escapement time-series. We also thank the fishery managers from CCFN who have contributed to discussions that lead to this paper through collaborative governance processes in the region.

## Author Contributions

KLW led the paper and all authors contributed to writing and editing. KLW, WIA, and BC led the analyses. KLW, MR, CNS, CKW, and WIA conceptualized the project. WIA and CKW led the data curation.

## Notes

https://doi.org/10.5281/zenodo.14714939

### Competing Interest Statement

The authors have declared no competing interest.

