## Appendix S1 for "Using fisheries risk assessment to inform precautionary and collaborative management in a declining coho salmon fishery"

**Supplemental S1. Trends in harvest rates**

Harvest rates for coho salmon are highly uncertain and remain unmonitored in most of the North and Central Coast, with the exception of three coded-wire tag (CWT) indicator stocks on the Nass (Zolzap Creek), Skeena (Toboggan Creek), and Haida Gwaii (Deena Creek). Given limited data, we made some simplifying assumptions about the magnitude of harvest in the NCC. First, we assumed that harvest rates of coho populations within the Skeena, Nass, and Haida Gwaii regions were equal to their local CWT indicator stocks. Second, we estimated exploitation rates for coho in regions that lacked CWT programs (PFMAs 5-10) based on modifications from previous work by English *et al.* (2018). For coho populations in PFMAs 5-10, English *et al.* (2018) estimated harvest rates based on Toboggan Creek and assumed that population-specific harvest rates from Canadian and Alaskan fisheries declined as a function of their location such that 100% of estimated harvest rates from Toboggan Creek were applied to more northern populations in PFMA 6, 60% for coho populations in PFMAs 5-9, and 40% of Canadian ER and 20% of Alaska for coho in PFMA 9-10. However, recent monitoring from the Central Coast First Nations suggests that total harvest for coho salmon is likely higher than initially assumed in English *et al.* (2018), with harvest estimates in the Central Coast region approximating that of PFMAs 3 and 4 (Steel *et al.* 2021). In addition, genetic stock identification suggests that Central Coast coho comprise ~20% of the total catch in commercial troll fisheries in Haida Gwaii (DFO *unpublished data*). We therefore assumed that populations in PFMAs 5-10 had higher harvest rates from Canadian fisheries than previously assumed and estimated domestic harvest rates for these regions by averaging the annual domestic harvest rates from the three CWT programs (Toboggan, Zolap, Deena). However, in the absence of data on Alaskan interceptions of populations without CWT marking and given the greater distances between PFMAs 5-10 and Alaskan coastal waters, we used Alaskan ERs estimated by English *et al.* (2018) following the assumptions detailed above.

Based on the above methods, coho harvest rates across the majority of NCC areas declined between 1980 and 2020, however the trend in harvest rates varied dramatically between CUs (Figure 2). All populations had higher harvest rates in the period from 1980 to 1997. From 1998 to 2003 harvest rates were dramatically reduced in Canadian domestic fisheries in response to a short-term collapse in coho populations during the early 1990s.

In general, harvest rates increased gradually after about 2002, and equaled or exceeded current levels by 2005. Coho returning to the Nass River, however, have been subjected to much higher harvest rates (2009-2019 mean = 52.5% SD 10.2%), and are disproportionately intercepted by Alaskan fisheries (mean Alaska ER = 35%, SD 13.8%). With the exception of the Nass River, recent harvest rates (2009-2019) have generally been moderate for coho populations in the NCC, ranging from an estimated low of 16.4% (SD 4%) for the Deena River CWT indicator, to a high of 38.2% (SD 5.8%) for Skeena River coho, and 38.1% (SD 4.4%) for coho in Area 6 rivers in Douglas Channel and the waters surrounding Kitimat. Estimated exploitation rates for Central Coast coho populations (PFMAs 5-10) are based on extrapolation from regional CWT indicators and should be interpreted with caution. However, harvest rates from 2009-2019 among the Central Coast regions were 21.8% for Rivers and Smiths Inlets, 25.1% for Hecate Strait Mainland, and 29.4% for other Central Coast populations in PFMAs 6-8.

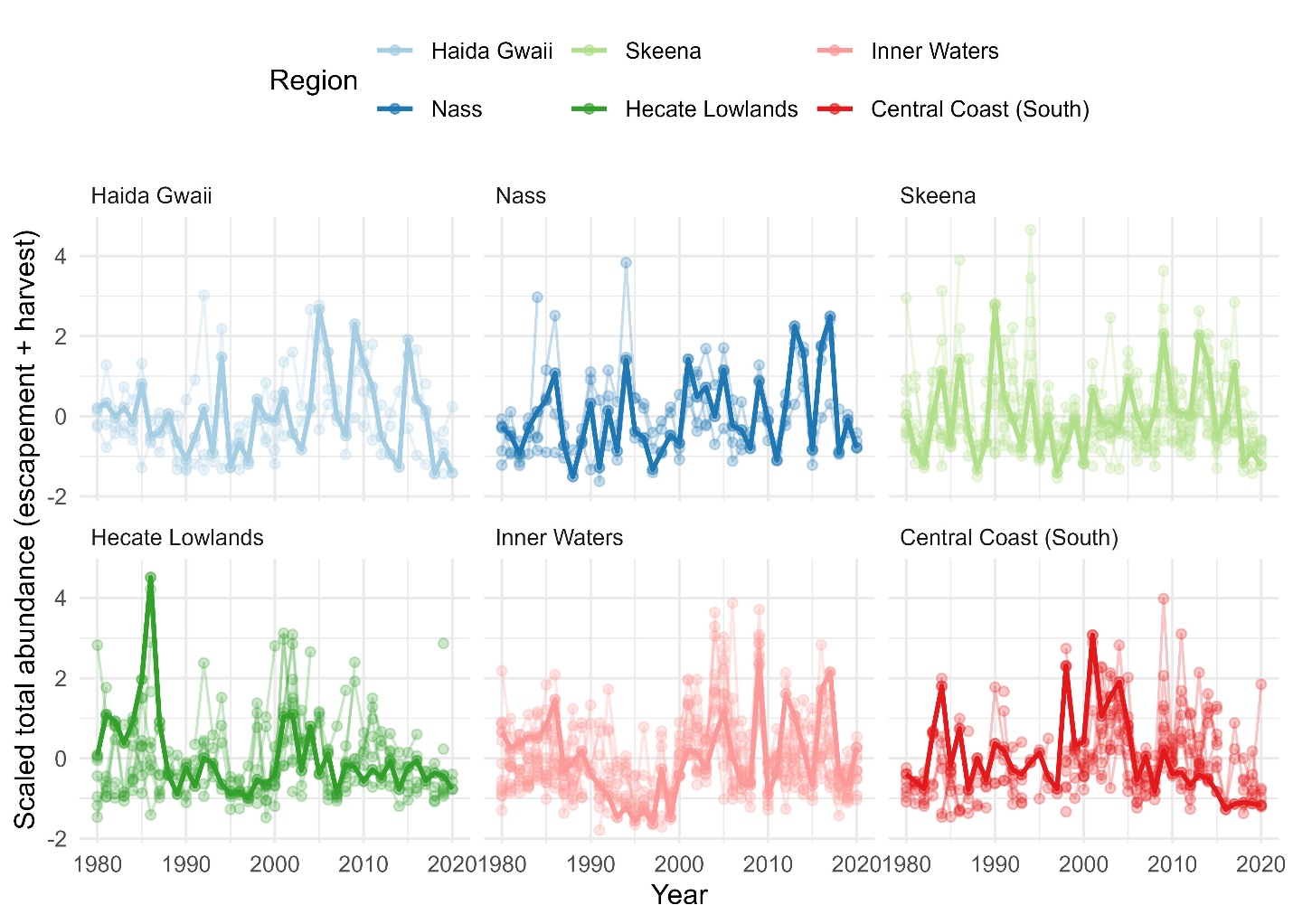

**Figure S1**. Total abundance (spawner escapement plus harvested salmon) of coho salmon since 1980. Abundance was mean centered and scaled by one standard deviation for each population within a region. Solid lines indicate the summed total abundance of coho populations within a region, which was then scale-transformed.

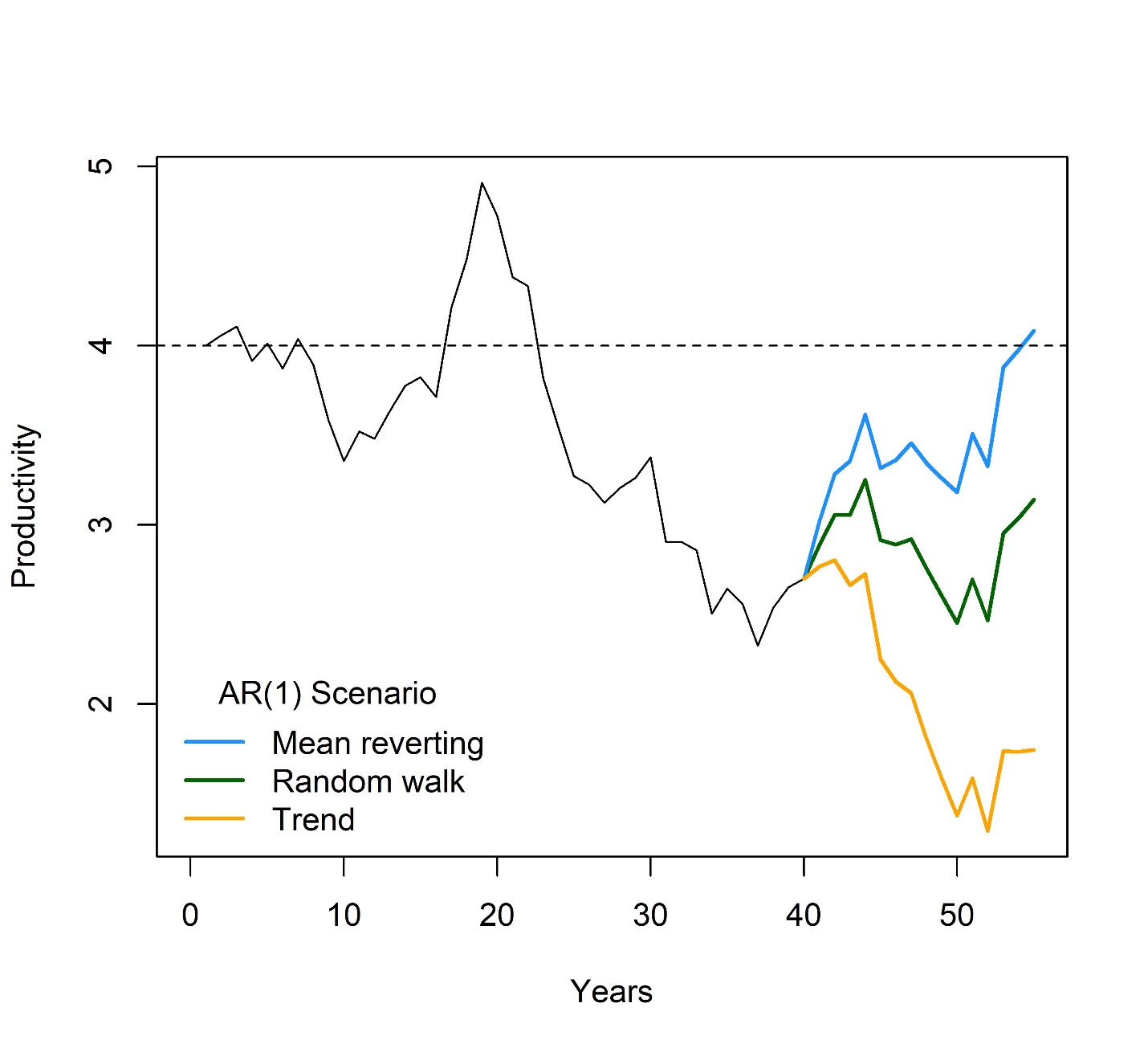

**Figure S2**. Example scenarios to model time-varying intrinsic productivity used during forward simulations of coho salmon population dynamics. As shown, productivity fluctuated as a random walk during years 1-20 and began a long-term decline in year 21-40. Beginning in year 41, three scenarios are demonstrated whereby time-varying productivity either: (1) reverted to a long-term mean of 4.0 (mean reverting), (2) followed a random walk based on the productivity from the previous year (darkgreen), or (3) continued on a long-term trend (yellow).

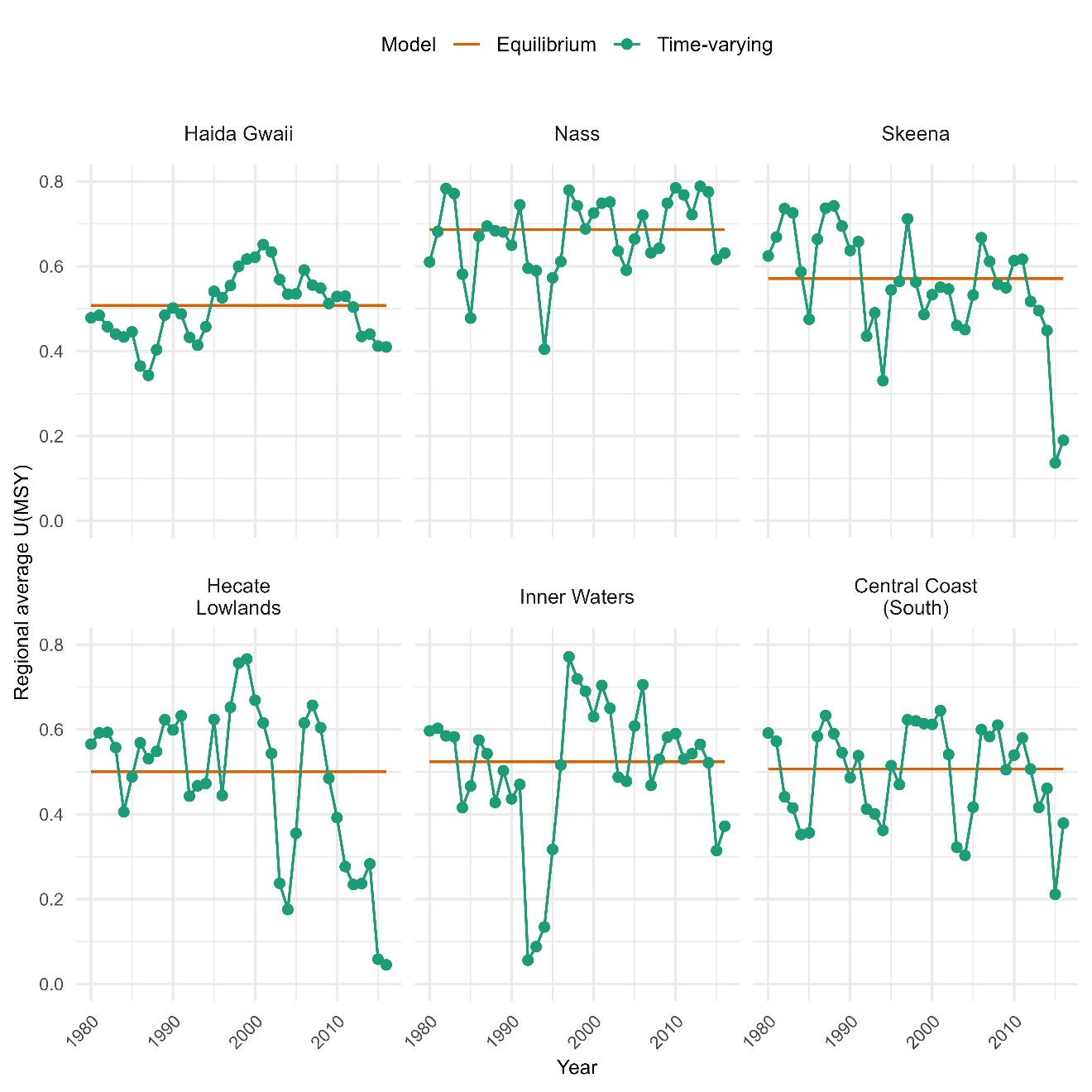

**Figure S3.** Posterior mean estimates of stationary (orange) and time-varying (green) UMSY reference point (harvest rate at maximum sustainable yield) among six regional groups of coho salmon populations along the North and Central Coast of British Columbia.

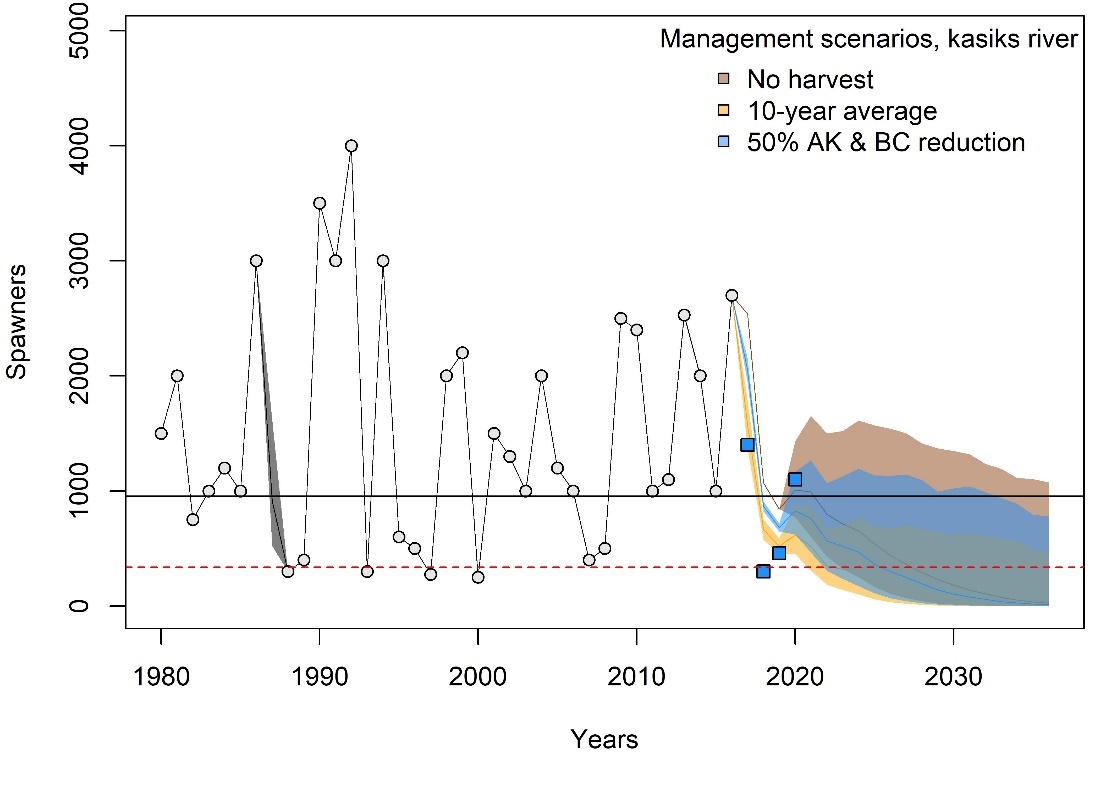

**Figure S4.** Coho salmon spawner escapement time-series from the Kasiks River and forward simulations under three management scenarios. Grey circles and lines (1980-2016) indicate observed escapement data used during model-fitting stage. The blue squares indicate observed escapement data from 2017-2020 used to evaluate predictive performance, while shaded polygons indicate inner 20% credible intervals (lines within shaded regions indicate posterior median estimates). Note, for ease of visuals we omitted showing two scenarios: (1) a 50% reduction in AK harvest and (2) a 50% reduction in BC harvest.

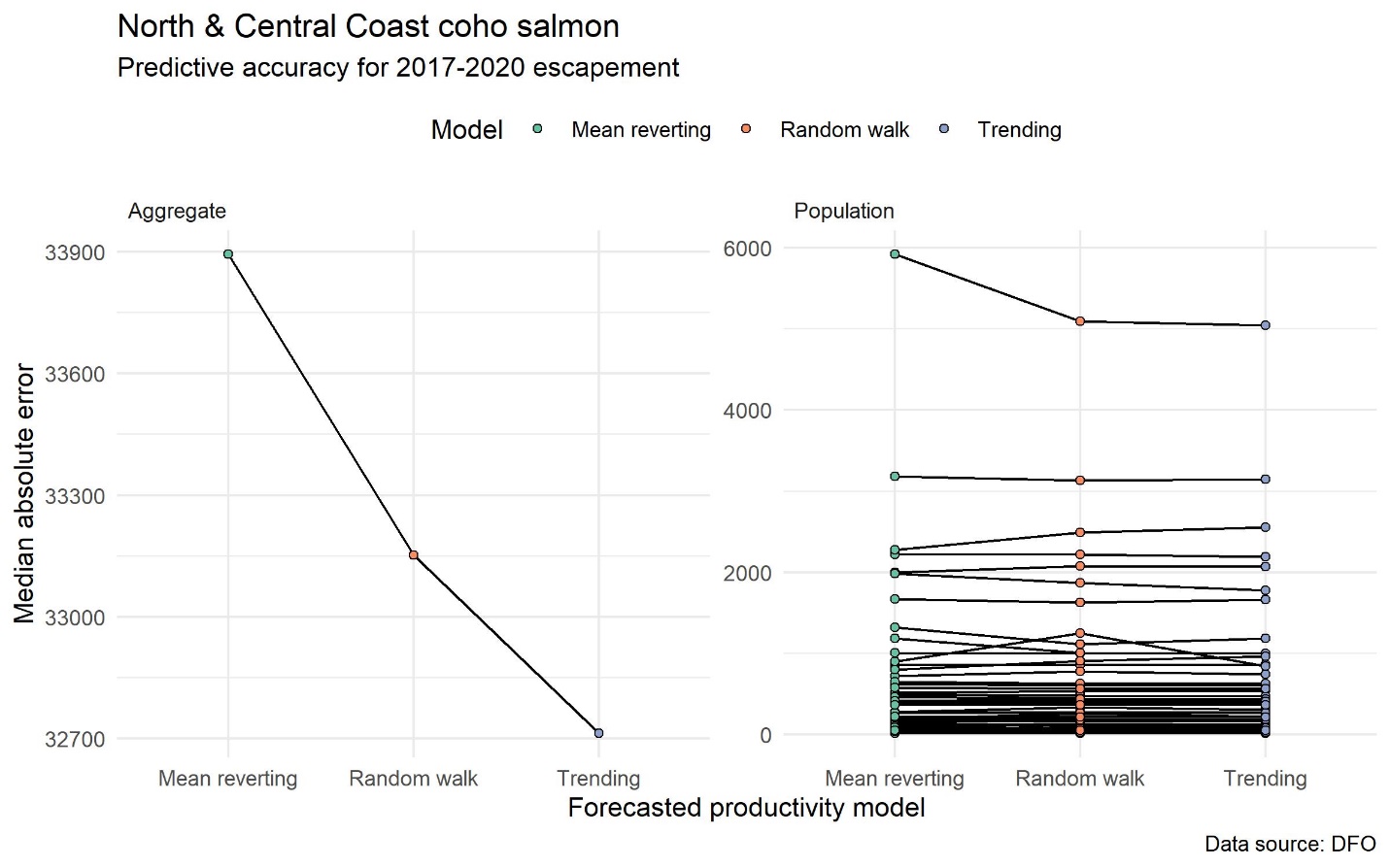

**Figure S5.** Median absolute error between observed and the posterior predictive distribution in spawner escapements from 2017-2020 at aggregate- (left) and population-levels (right) to contrast three different forward simulation models: **mean reverting** (green points), where productivity was expected to return to a long-term average; **random walk** (orange points), where productivity continued forward on a correlated random walk; and **trending (**blue points), where productivity continued forward along a long-term trend. Lower values indicate higher predictive accuracy. Note that the 2017-2020 data was not used in training the estimated productivity trends and thus were a test dataset in a limited cross-validation exercise.

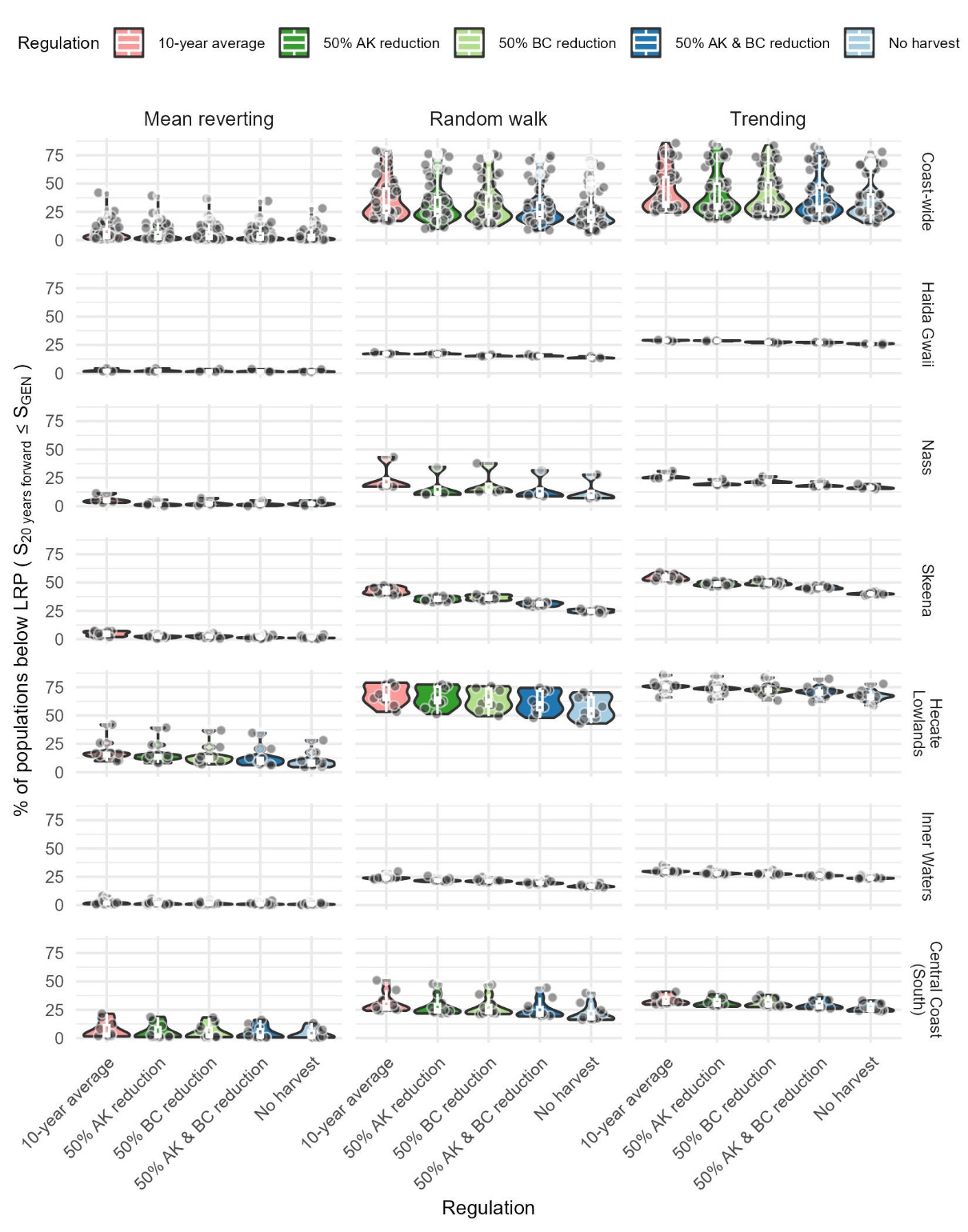

**Figure S6.** Posterior frequency distributions that future years of North and Central Coast coho salmon populations (points) fell below the limit reference point (LRP) under forward simulations of three productivity and five harvest management scenarios at aggregate (top panel) and regional scales.

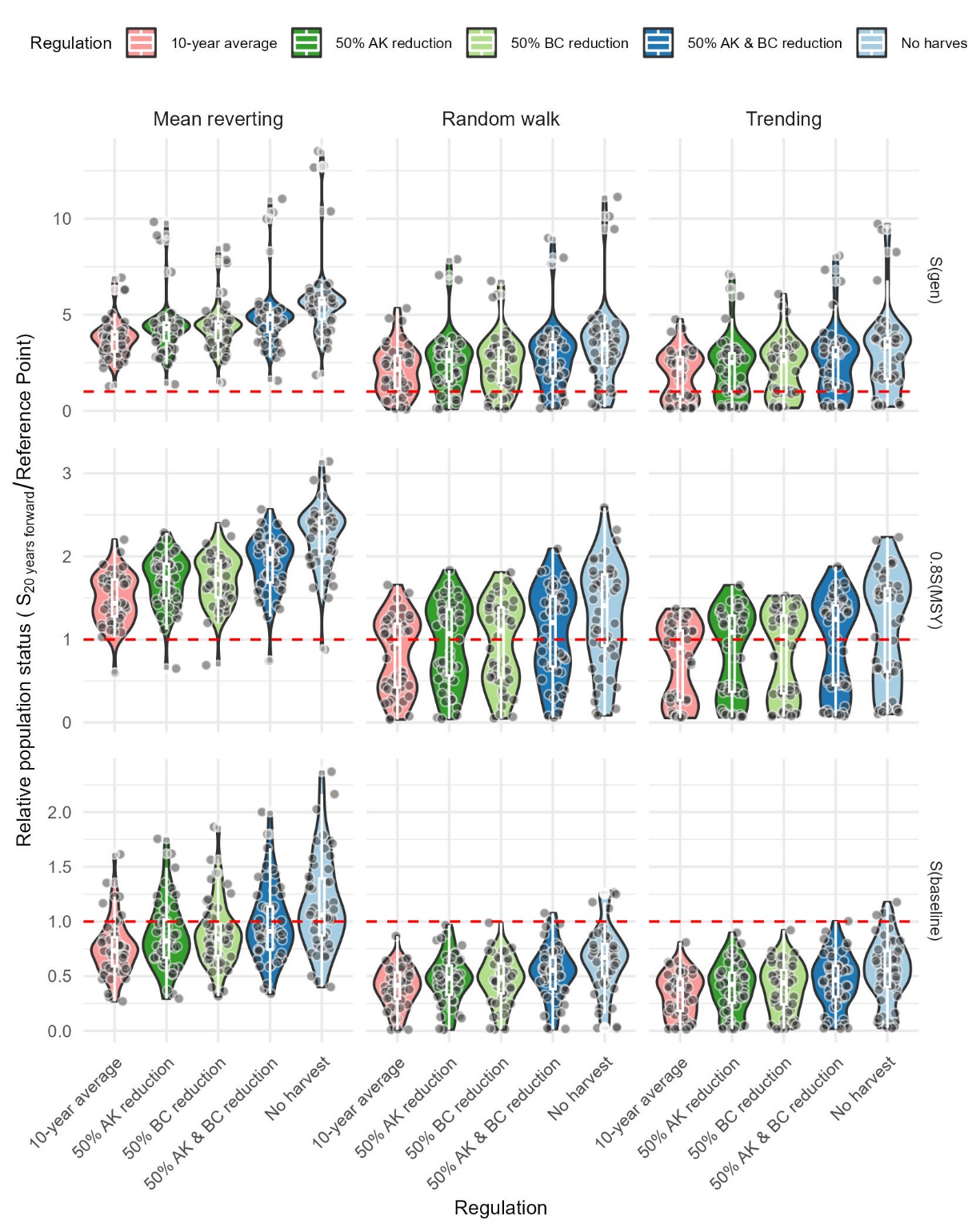

**Figure S7.** Posterior distributions of the relative population status of North and Central Coast coho salmon populations under forward simulations of three productivity and five harvest management scenarios at the aggregate, coastal scale.

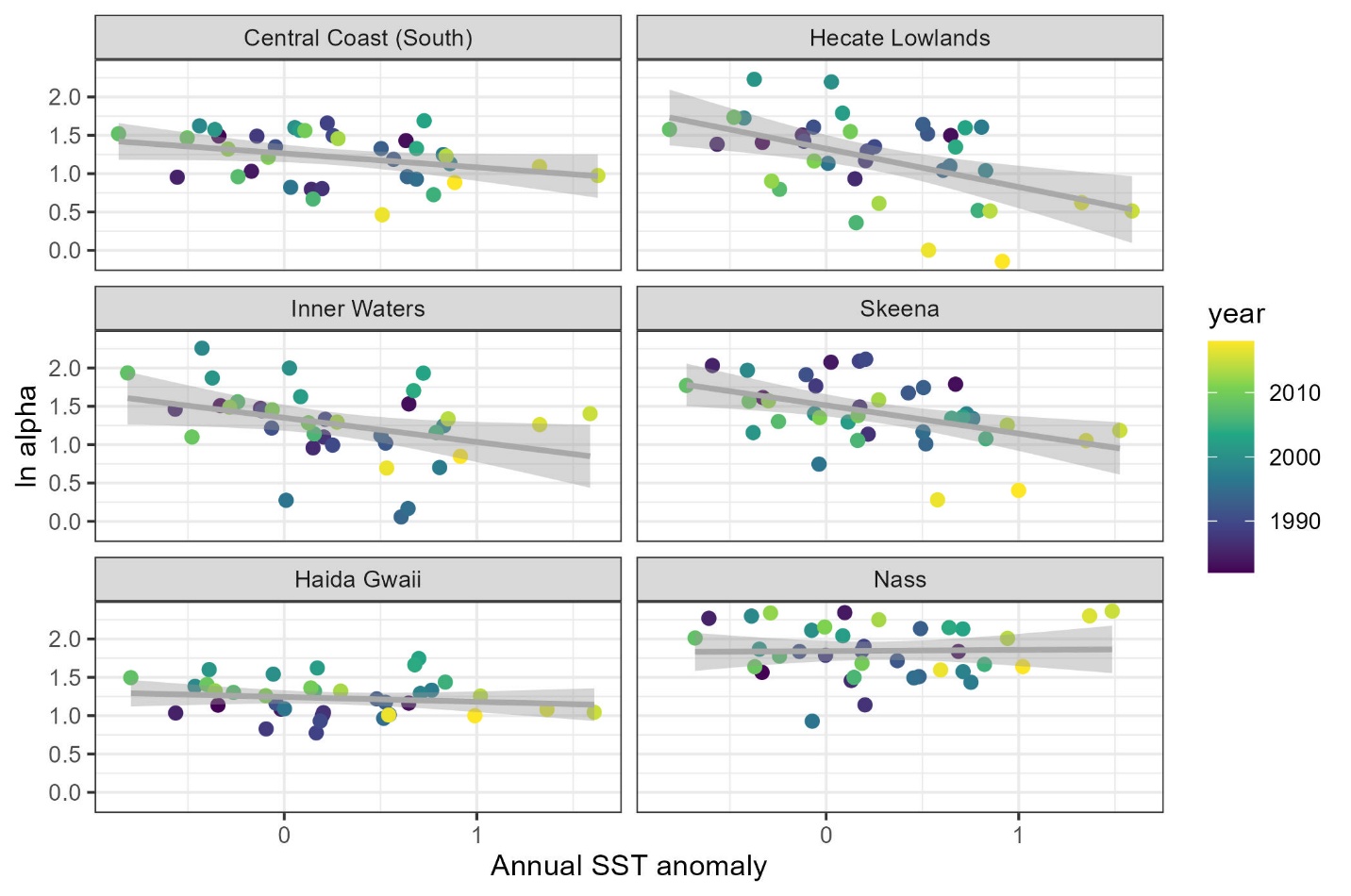

**Figure S8:** Relationships between annual sea surface temperature anomalies and regional productivity values for NCC coho salmon populations. Temperature data were lagged two years to reflect the year of ocean entry.

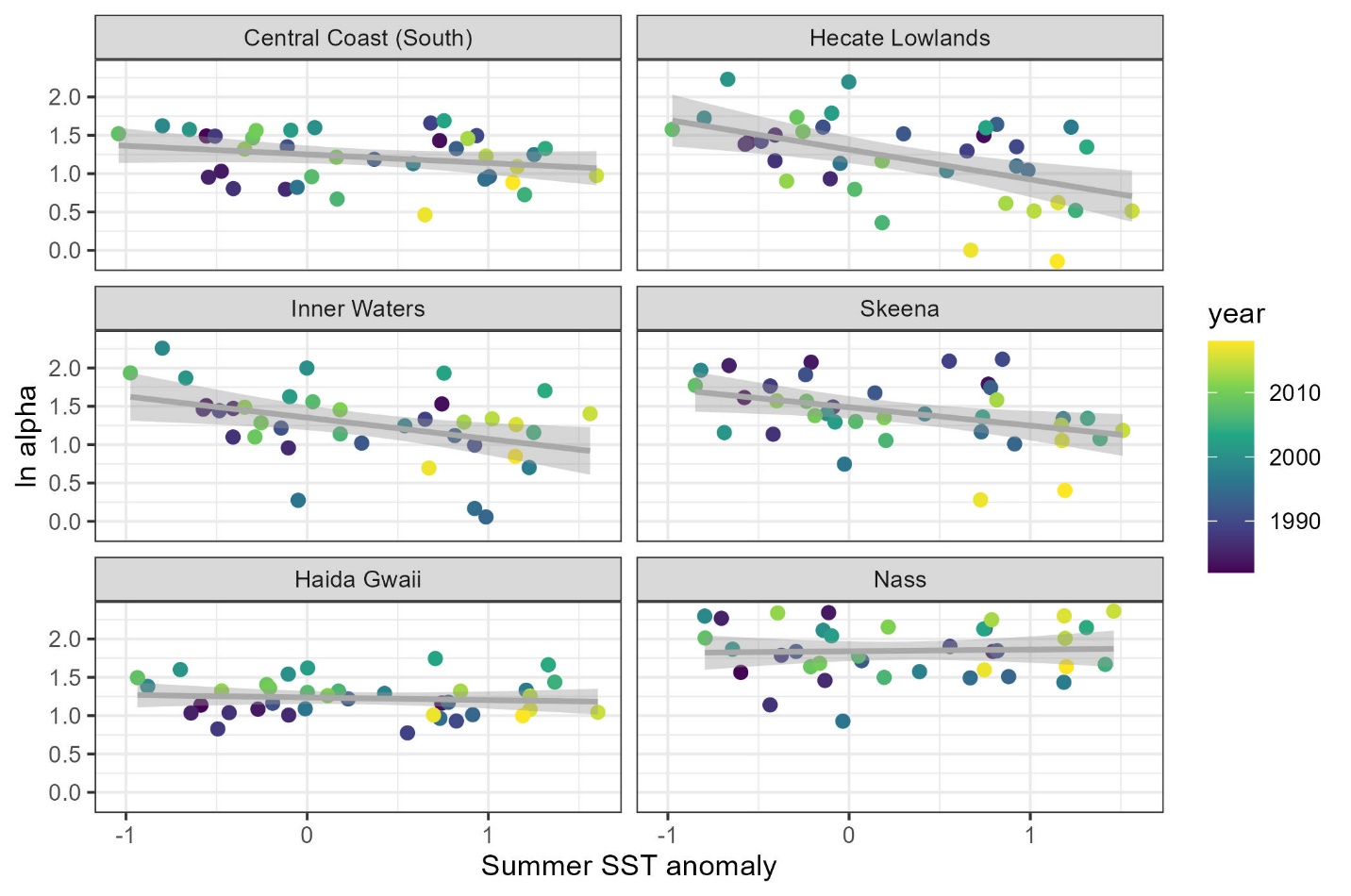

**Figure S9:** Relationships between summer sea surface temperature anomalies and regional productivity values for NCC coho salmon populations. Temperature data were lagged two years to reflect the year of ocean entry.

| Table S1 – The average adult proportion-at-age, combining duration of freshwater and at-sea rearing, for coho salmon populations across North and Central Coast of British Columbia since 1980. | | | |
| --- | --- | --- | --- |
| Population | Age-3 | Age-4 | Age-5 |
| Aaltanhash | 0.714 | 0.286 | 0.000 |
| Arnoup | 0.854 | 0.141 | 0.004 |
| Babine Fence | 0.752 | 0.246 | 0.002 |
| Bear | 0.615 | 0.380 | 0.005 |
| Bella Coola | 0.946 | 0.054 | 0.000 |
| Belowe Creek | 0.854 | 0.141 | 0.004 |
| Brim | 0.714 | 0.286 | 0.000 |
| Cascade | 0.714 | 0.286 | 0.000 |
| Chuckwalla | 0.794 | 0.206 | 0.000 |
| Dala | 0.714 | 0.286 | 0.000 |
| Damshilgwit | 0.615 | 0.380 | 0.005 |
| Deena | 0.854 | 0.141 | 0.004 |
| Diskangieg | 0.571 | 0.426 | 0.003 |
| Docee | 0.775 | 0.225 | 0.000 |
| East Arm | 0.854 | 0.141 | 0.004 |
| Ecstall | 0.571 | 0.426 | 0.003 |
| Elcho | 0.714 | 0.286 | 0.000 |
| Evelyn Creek | 0.714 | 0.286 | 0.000 |
| Exchamsiks | 0.571 | 0.426 | 0.003 |
| Exstew | 0.571 | 0.426 | 0.003 |
| Foch Creek | 0.714 | 0.286 | 0.000 |
| Green | 0.714 | 0.286 | 0.000 |
| Hartley Bay Creek | 0.854 | 0.141 | 0.004 |
| Hugh Ck | 0.714 | 0.286 | 0.000 |
| Kadeen | 0.571 | 0.426 | 0.003 |
| Kasiks | 0.571 | 0.426 | 0.003 |
| Kemano | 0.714 | 0.286 | 0.000 |
| Kildala | 0.714 | 0.286 | 0.000 |
| Kiltush | 0.714 | 0.286 | 0.000 |
| Kiskosh Creek | 0.714 | 0.286 | 0.000 |
| Kitwanga | 0.752 | 0.246 | 0.002 |
| Lachmach | 0.548 | 0.450 | 0.003 |
| Martin | 0.714 | 0.286 | 0.000 |
| Meziadin | 0.681 | 0.318 | 0.001 |
| Nangeese kspx | 0.752 | 0.246 | 0.002 |
| Necleetsconnay | 0.946 | 0.054 | 0.000 |
| Nias | 0.854 | 0.141 | 0.004 |
| Pallant Creek | 0.854 | 0.141 | 0.004 |
| Paril | 0.714 | 0.286 | 0.000 |
| Quaal | 0.854 | 0.141 | 0.004 |
| Quartcha | 0.714 | 0.286 | 0.000 |
| Riordan | 0.714 | 0.286 | 0.000 |
| Roscoe | 0.714 | 0.286 | 0.000 |
| Salloomt | 0.946 | 0.054 | 0.000 |
| Sylvia Creek | 0.854 | 0.141 | 0.004 |
| Tlell | 0.854 | 0.141 | 0.004 |
| Tsimtack Lake | 0.854 | 0.141 | 0.004 |
| Tyler Creek | 0.854 | 0.141 | 0.004 |
| Wahoo | 0.714 | 0.286 | 0.000 |
| West Arm | 0.854 | 0.141 | 0.004 |
| Zolzap | 0.571 | 0.426 | 0.003 |
| Zymagotitz | 0.571 | 0.426 | 0.003 |
