## Supplemental Data 3 for "Using fisheries risk assessment to inform precautionary and collaborative management in a declining coho salmon fishery"

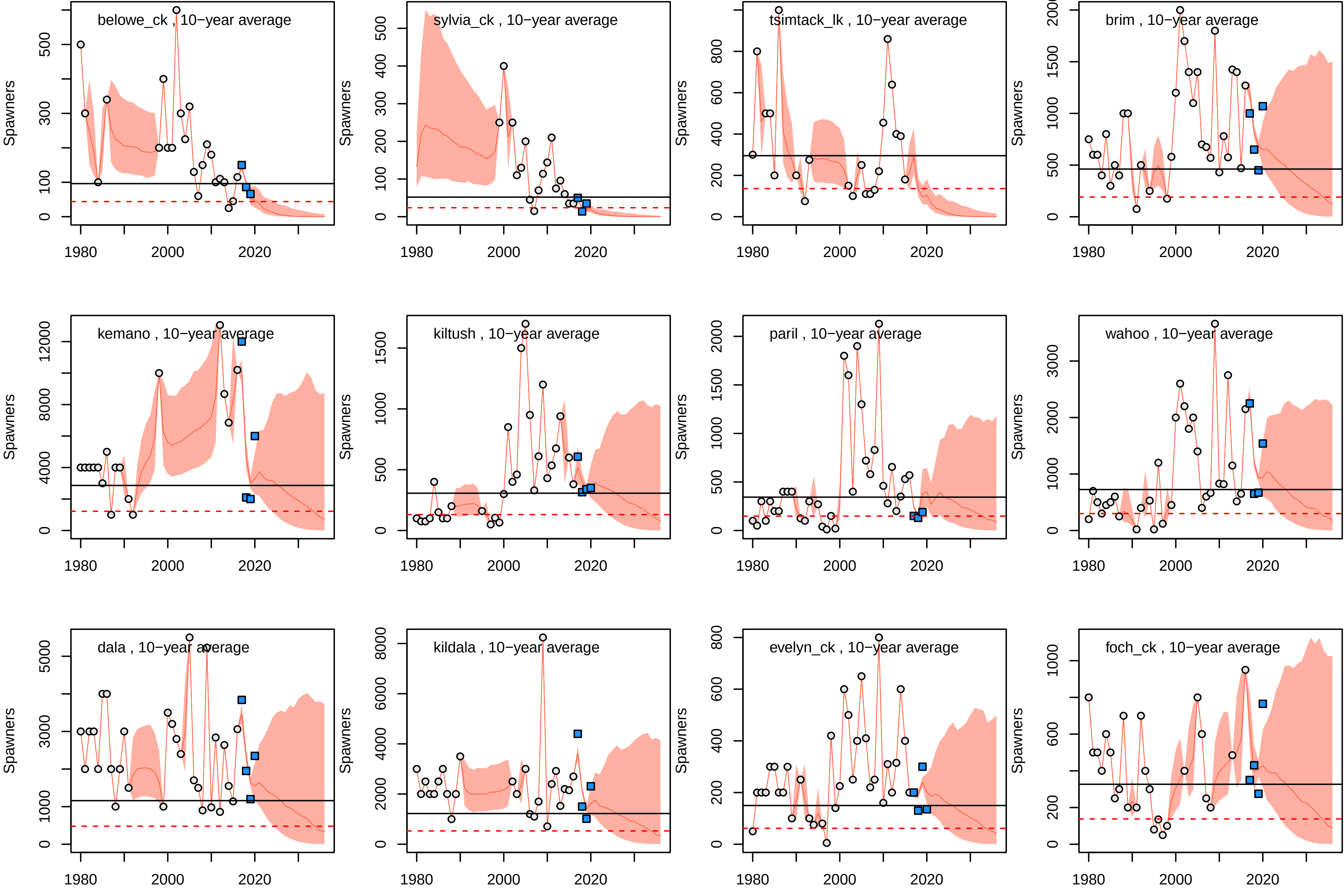
belowe_ck , 10−year average
sylvia_ck , 10−year average
tsimtack_lk , 10−year average
brim , 10−year average
kemano , 10−year average
kiltush , 10−year average
paril , 10−year average
wahoo , 10−year average
dala , 10−year average
kildala , 10−year average
evelyn_ck , 10−year average
foch_ck , 10−year average
Spawners
1980
2000
2020

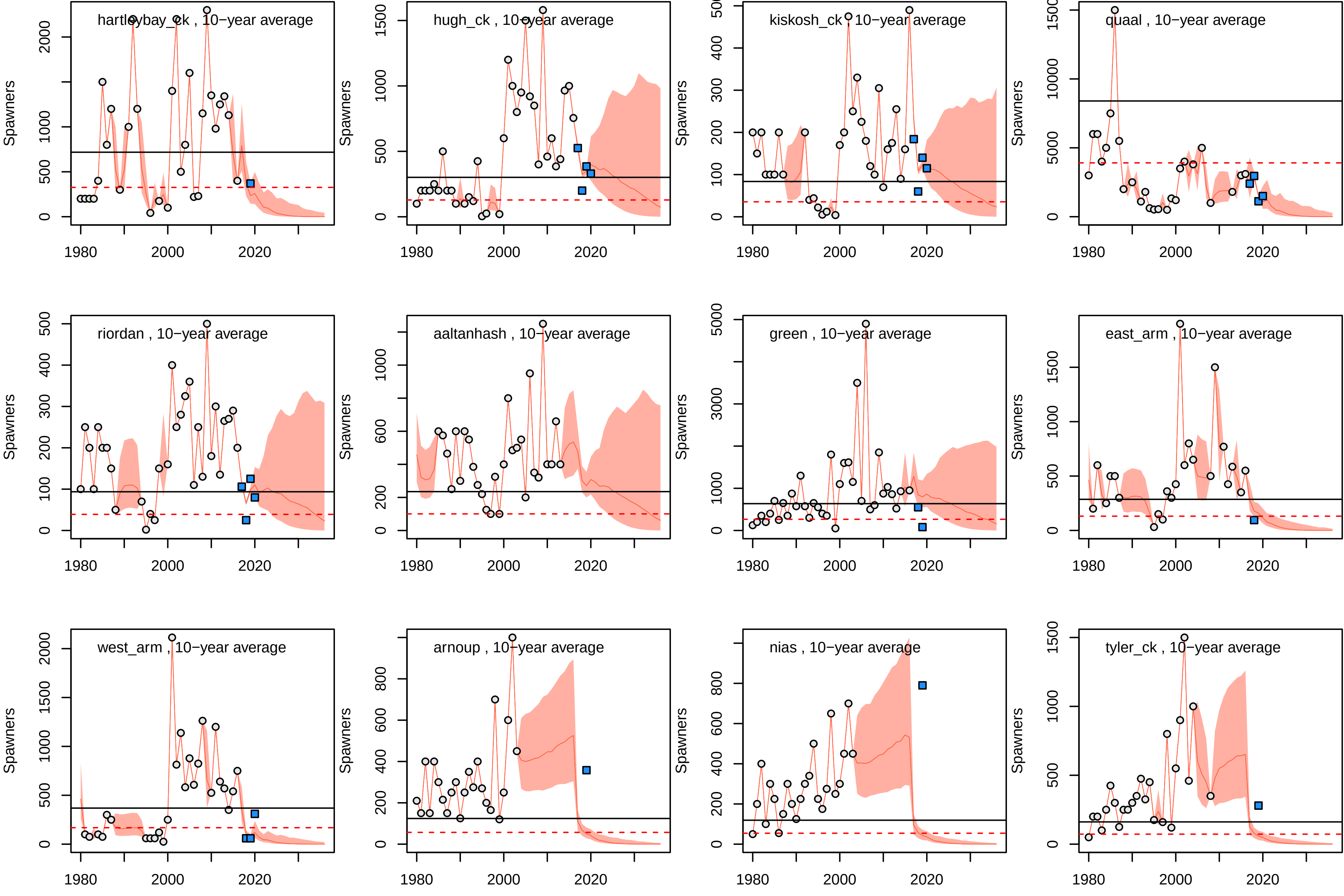
hartleybay_ck , 10−year average
hugh_ck , 10−year average
kiskosh_ck , 10−year average
quaal , 10−year average
riordan , 10−year average
aaltanhash , 10−year average
green , 10−year average
east_arm , 10−year average
west_arm , 10−year average
arnoup , 10−year average
nias , 10−year average
tyler_ck , 10−year average
Spawners
1980
2000
2020

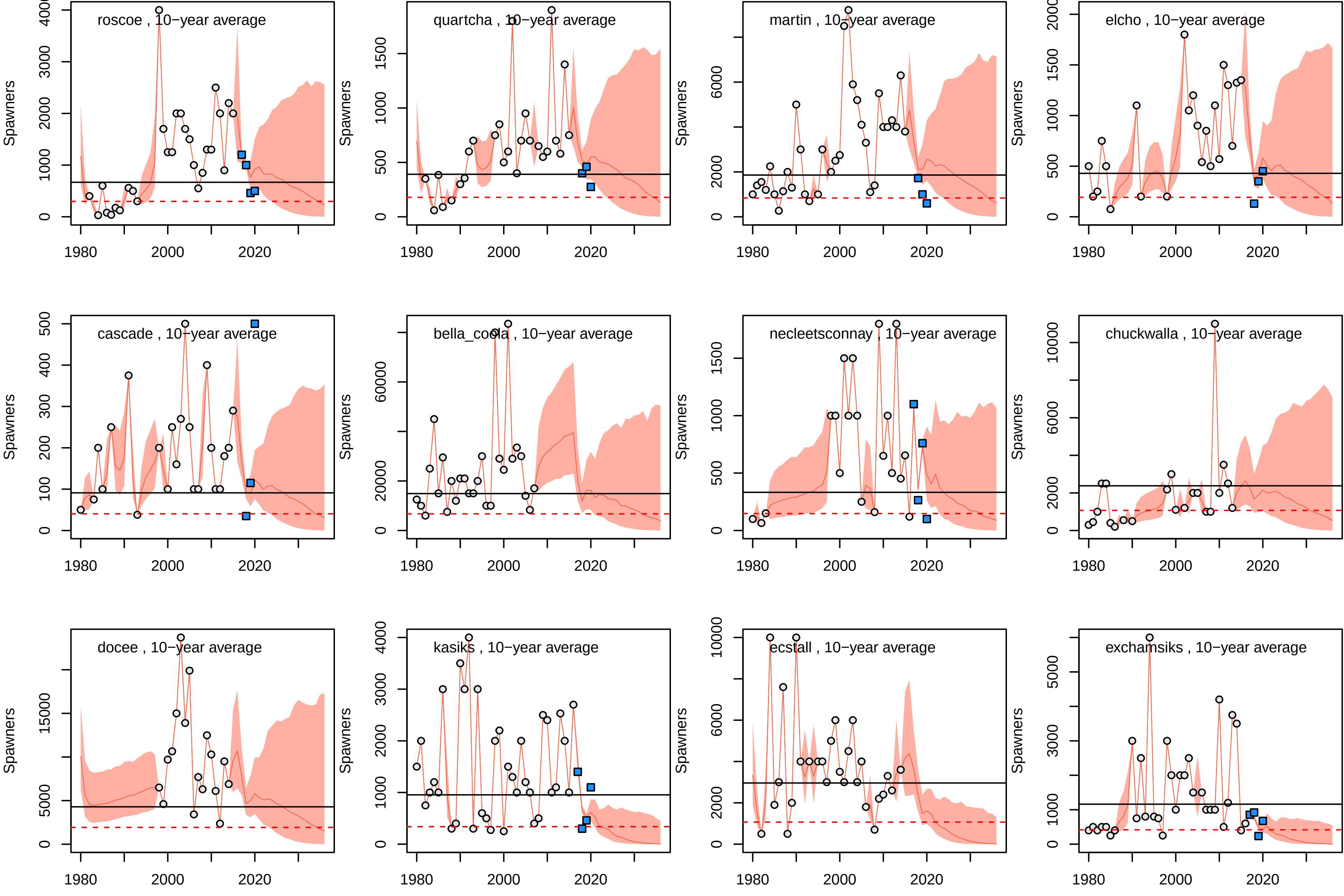
roscoe , 10−year average
quartcha , 10−year average
martin , 10−year average
elcho , 10−year average
cascade , 10−year average
bella_coola , 10−year average
necleetsconnay , 10−year average
chuckwalla , 10−year average
docee , 10−year average
kasiks , 10−year average
ecstall , 10−year average
exchamsiks , 10−year average
Spawners
1980
2000
2020

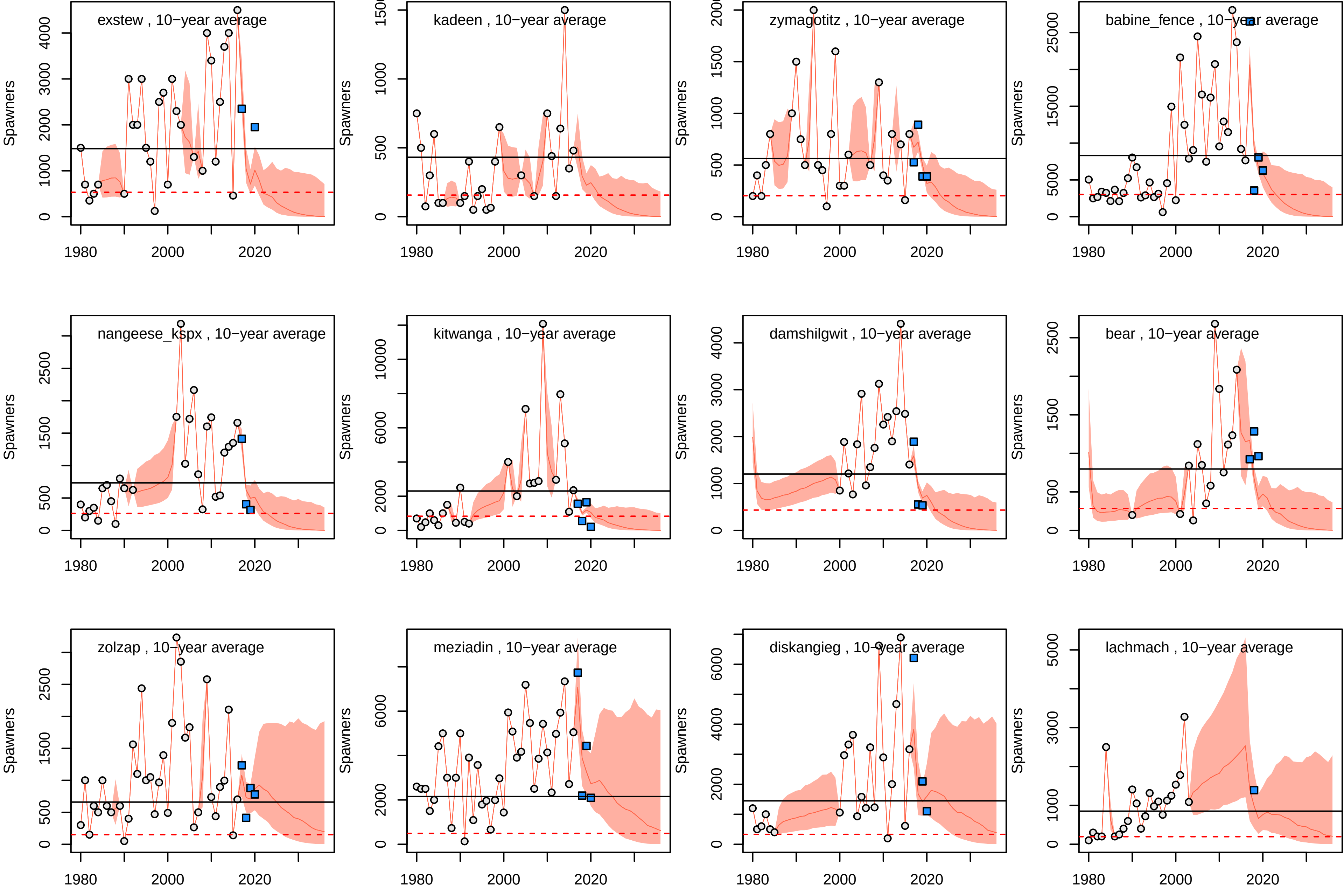
exstew , 10−year average
Spawners
0
1000
2000
3000
4000
1980
2000
2020
kadeen , 10−year average
Spawners
0
500
1000
1500
1980
2000
2020
zymagotitz , 10−year average
Spawners
0
500
1000
1500
2000
1980
2000
2020
babine_fence , 10−year average
Spawners
0
5000
15000
25000
1980
2000
2020
nangeese_kspx , 10−year average
Spawners
0
500
1500
2500
1980
2000
2020
kitwanga , 10−year average
Spawners
0
2000
6000
10000
1980
2000
2020
damshilgwit , 10−year average
Spawners
0
1000
2000
3000
4000
1980
2000
2020
bear , 10−year average
Spawners
0
500
1500
2500
1980
2000
2020
zolzap , 10−year average
Spawners
0
500
1500
2500
1980
2000
2020
meziadin , 10−year average
Spawners
0
2000
6000
1980
2000
2020
diskangieg , 10−year average
Spawners
0
2000
4000
6000
1980
2000
2020
lachmach , 10−year average
Spawners
0
1000
3000
5000
1980
2000
2020

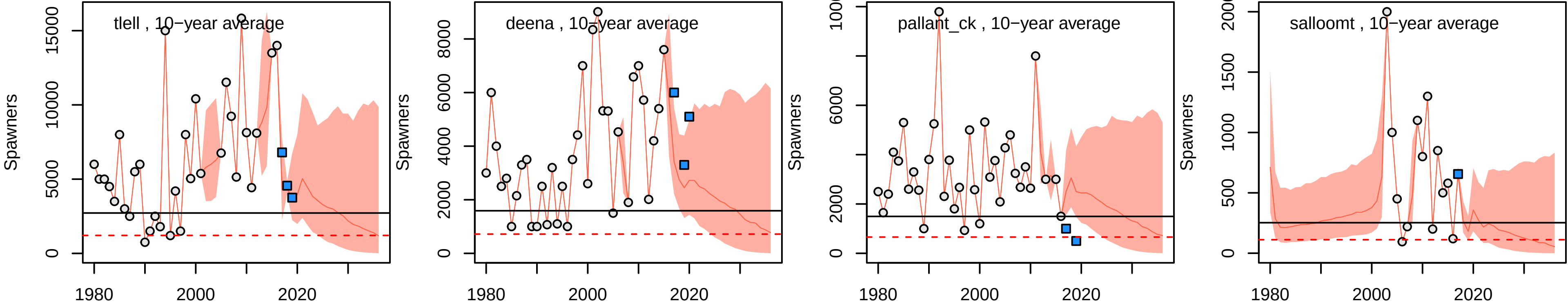
tlell , 10−year average
deena , 10−year average
pallant_ck , 10−year average
salloomt , 10−year average
Spawners
1980
2000
2020

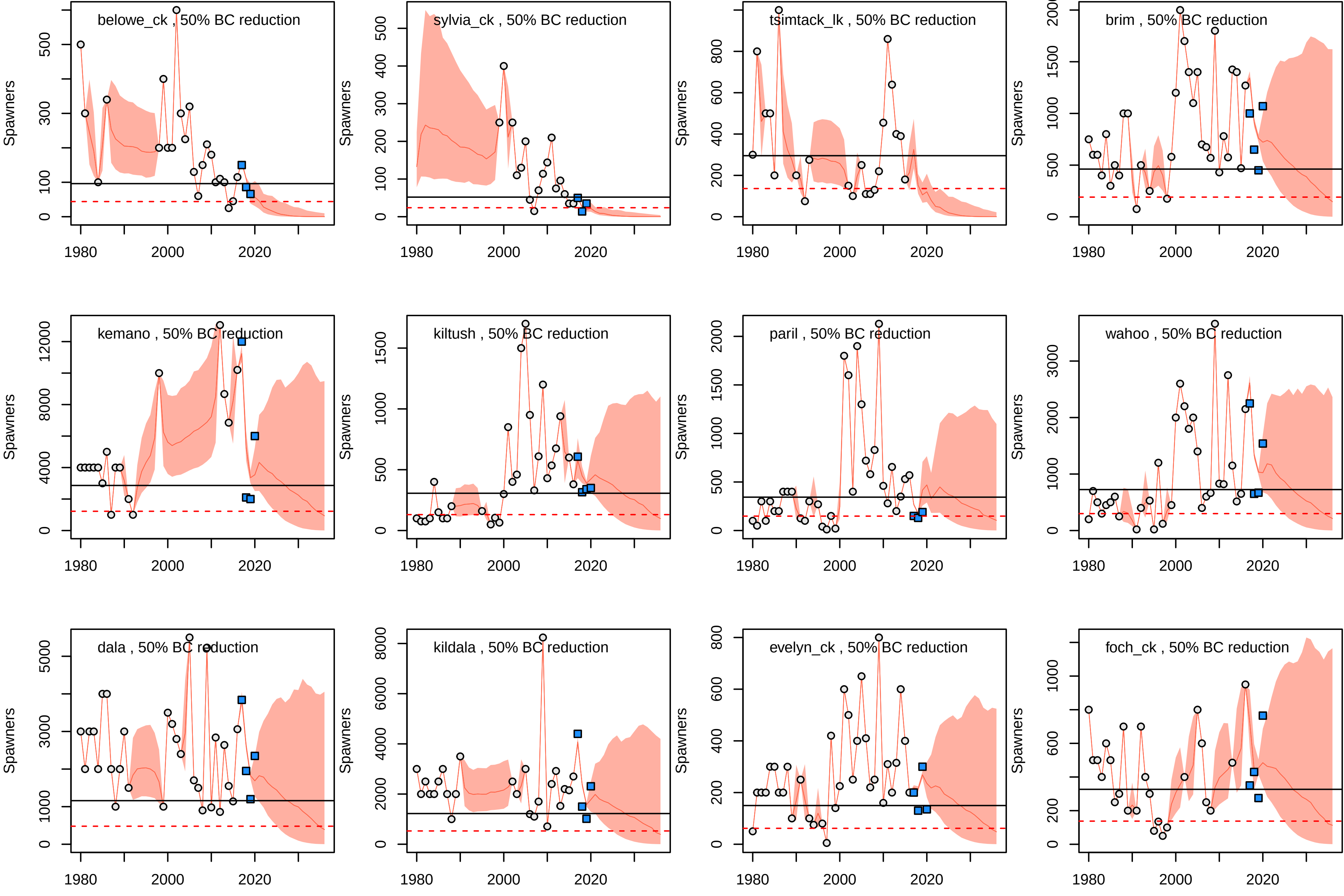
belowe_ck , 50% BC reduction
sylvia_ck , 50% BC reduction
tsimtack_lk , 50% BC reduction
brim , 50% BC reduction
kemano , 50% BC reduction
kiltush , 50% BC reduction
paril , 50% BC reduction
wahoo , 50% BC reduction
dala , 50% BC reduction
kildala , 50% BC reduction
evelyn_ck , 50% BC reduction
foch_ck , 50% BC reduction
Spawners
1980
2000
2020

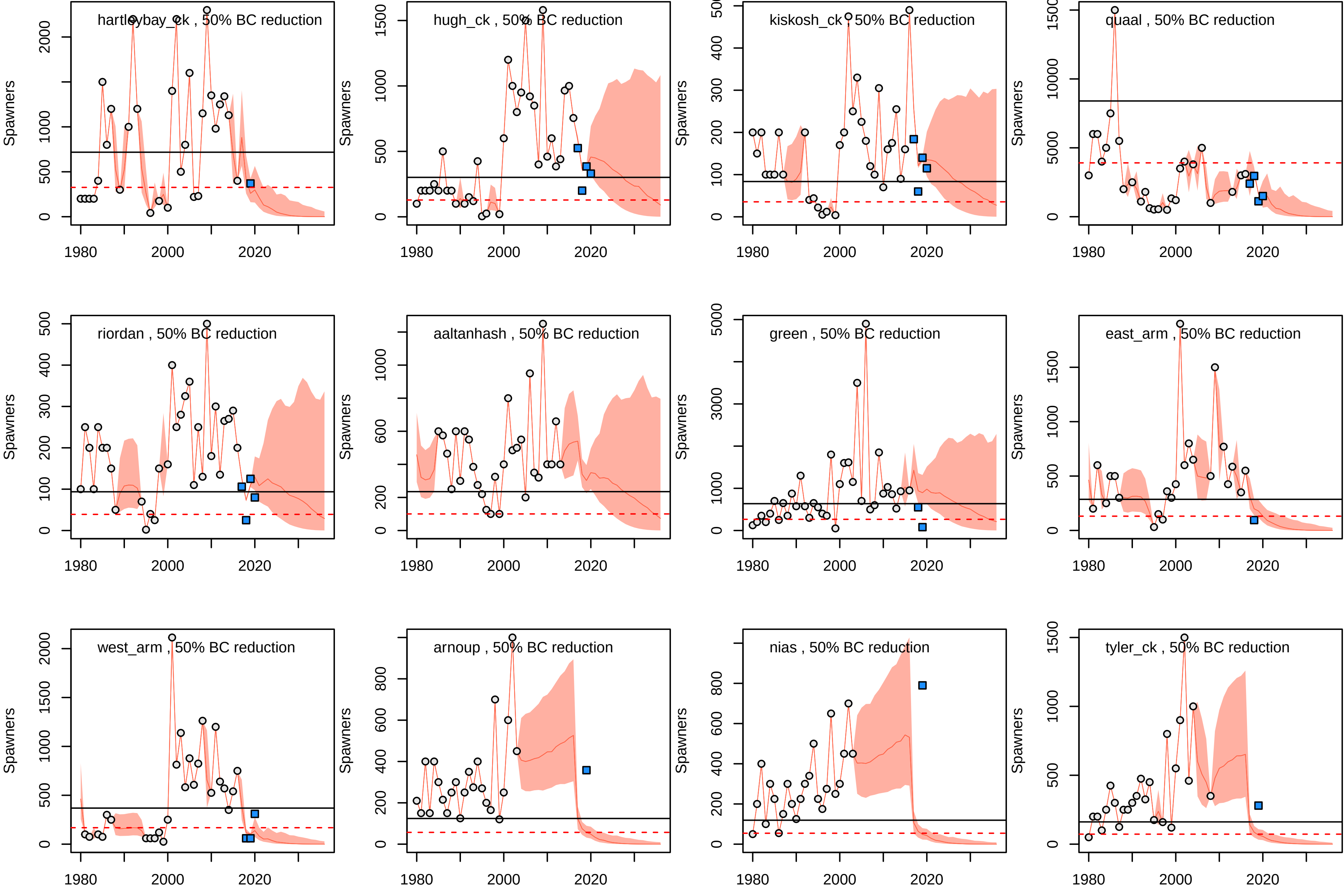
hartleybay_ck , 50% BC reduction
hugh_ck , 50% BC reduction
kiskosh_ck , 50% BC reduction
quaal , 50% BC reduction
riordan , 50% BC reduction
aaltanhash , 50% BC reduction
green , 50% BC reduction
east_arm , 50% BC reduction
west_arm , 50% BC reduction
arnoup , 50% BC reduction
nias , 50% BC reduction
tyler_ck , 50% BC reduction
Spawners
1980
2000
2020

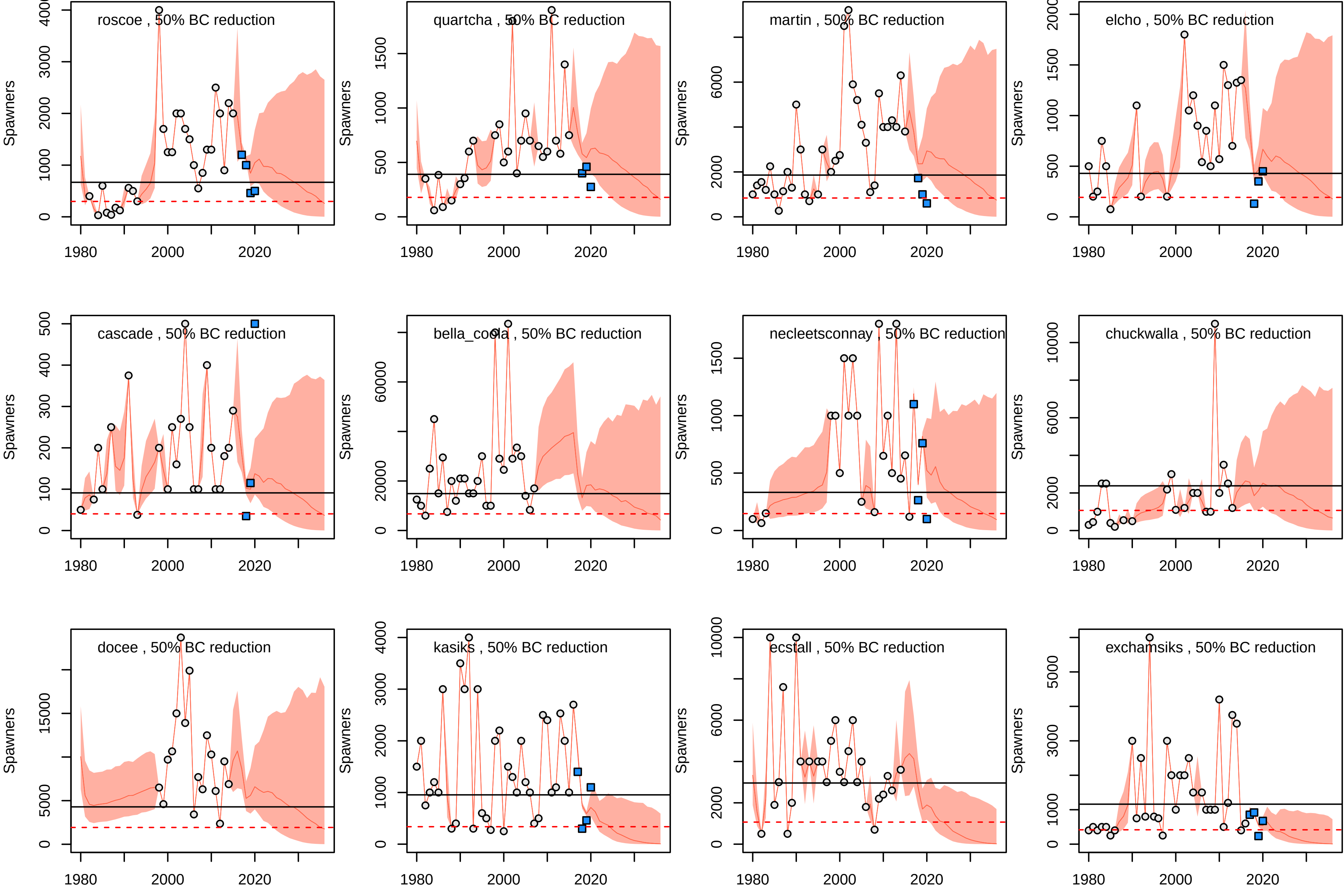
roscoe , 50% BC reduction
quartcha , 50% BC reduction
martin , 50% BC reduction
elcho , 50% BC reduction
cascade , 50% BC reduction
bella_coola , 50% BC reduction
necleetsconnay , 50% BC reduction
chuckwalla , 50% BC reduction
docee , 50% BC reduction
kasiks , 50% BC reduction
ecstall , 50% BC reduction
exchamsiks , 50% BC reduction
Spawners
1980
2000
2020

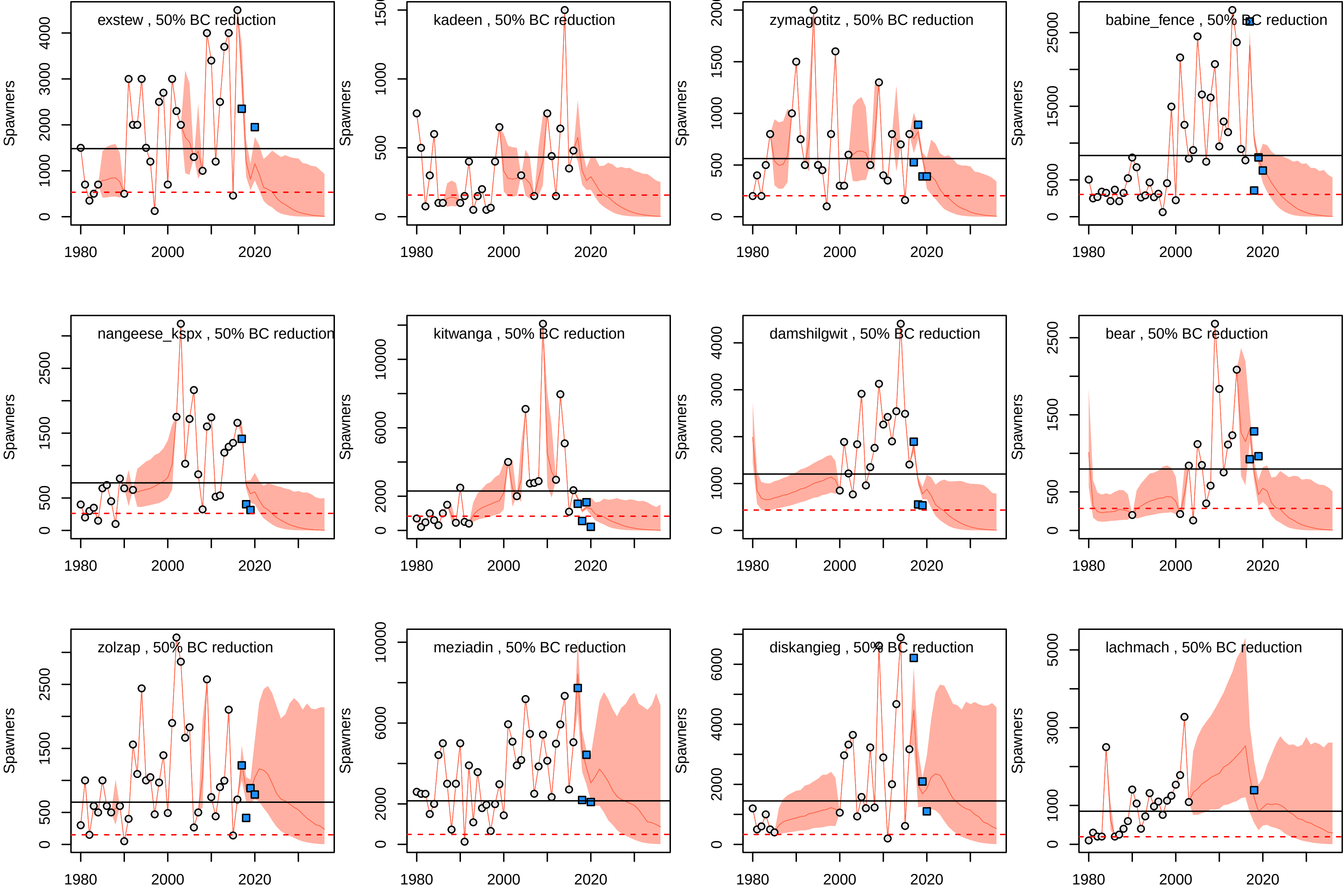
exstew , 50% BC reduction
kadeen , 50% BC reduction
zymagotitz , 50% BC reduction
babine_fence , 50% BC reduction
nangeese_kspx , 50% BC reduction
kitwanga , 50% BC reduction
damshilgwit , 50% BC reduction
bear , 50% BC reduction
zolzap , 50% BC reduction
meziadin , 50% BC reduction
diskangieg , 50% BC reduction
lachmach , 50% BC reduction
Spawners
1980
2000
2020

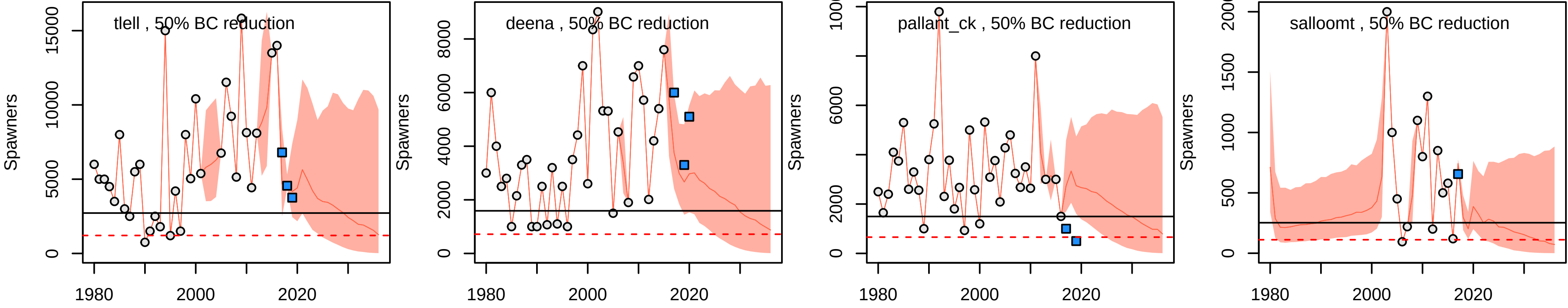

tlell , 50% BC reduction
deena , 50% BC reduction
pallant_ck , 50% BC reduction
salloomt , 50% BC reduction
Spawners
1980
2000
2020

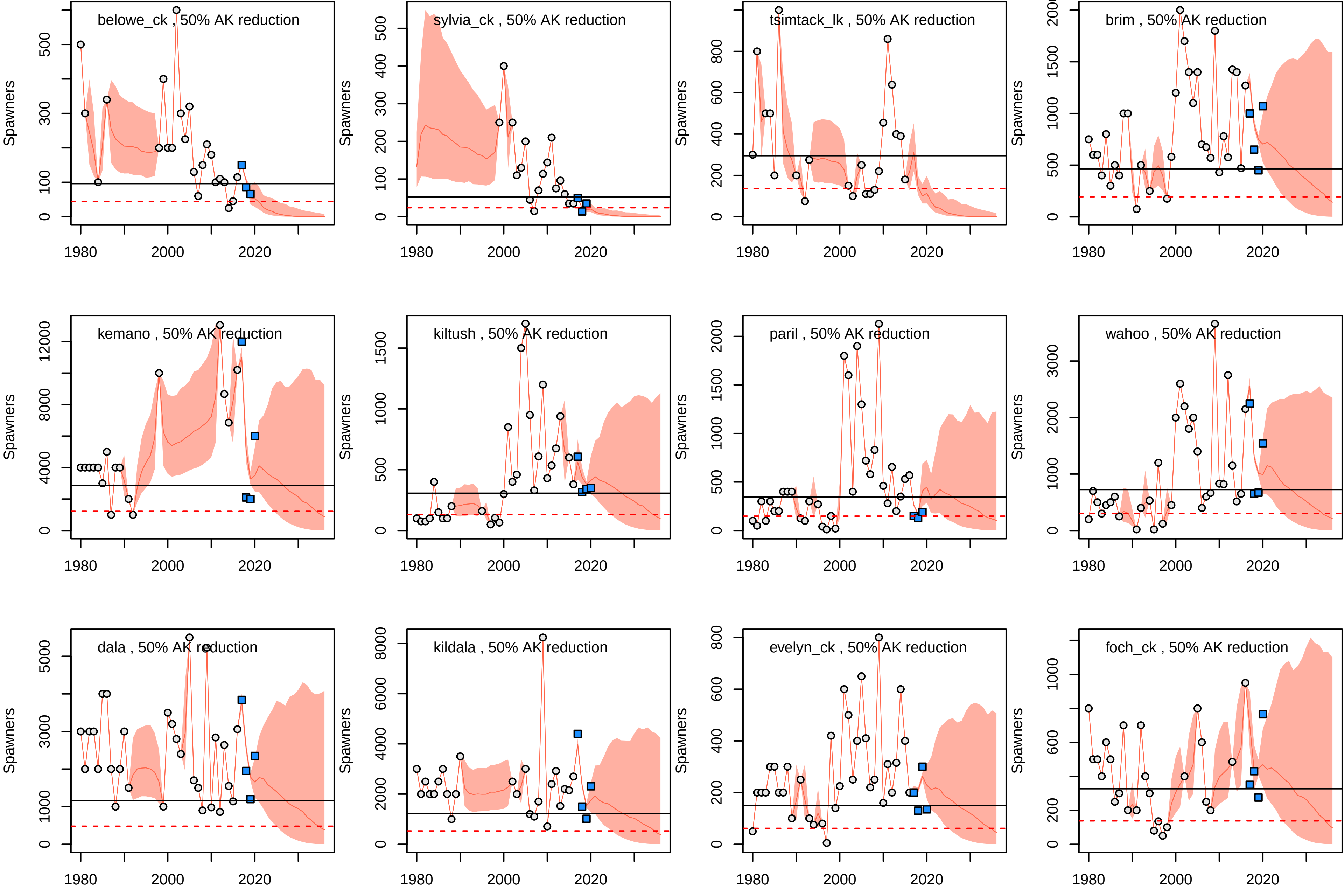

belowe_ck , 50% AK reduction
sylvia_ck , 50% AK reduction
tsimtack_lk , 50% AK reduction
brim , 50% AK reduction
kemano , 50% AK reduction
kiltush , 50% AK reduction
paril , 50% AK reduction
wahoo , 50% AK reduction
dala , 50% AK reduction
kildala , 50% AK reduction
evelyn_ck , 50% AK reduction
foch_ck , 50% AK reduction
Spawners
1980
2000
2020

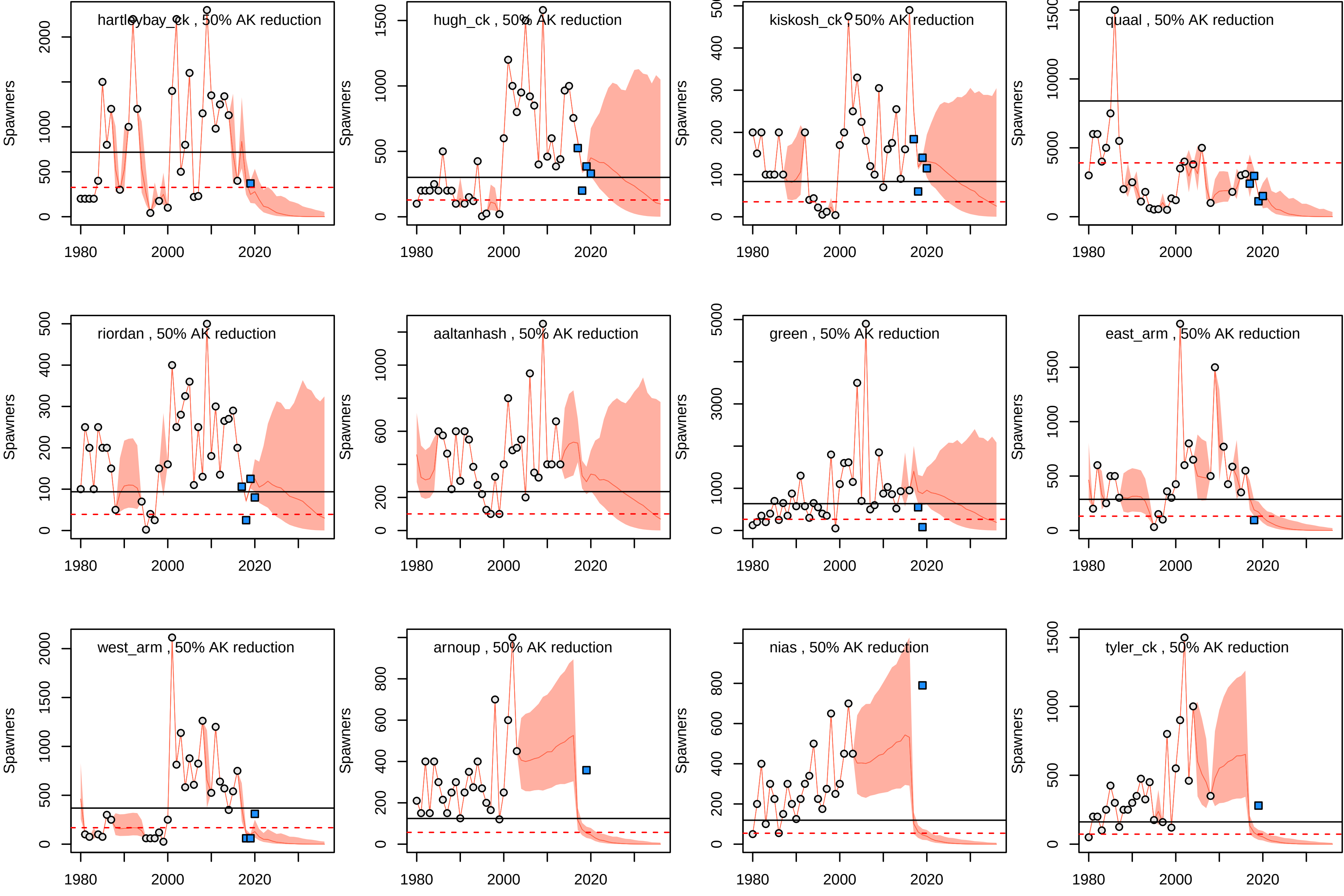
hartleybay_ck , 50% AK reduction
hugh_ck , 50% AK reduction
kiskosh_ck , 50% AK reduction
quaal , 50% AK reduction
riordan , 50% AK reduction
aaltanhash , 50% AK reduction
green , 50% AK reduction
east_arm , 50% AK reduction
west_arm , 50% AK reduction
arnoup , 50% AK reduction
nias , 50% AK reduction
tyler_ck , 50% AK reduction
Spawners
1980
2000
2020

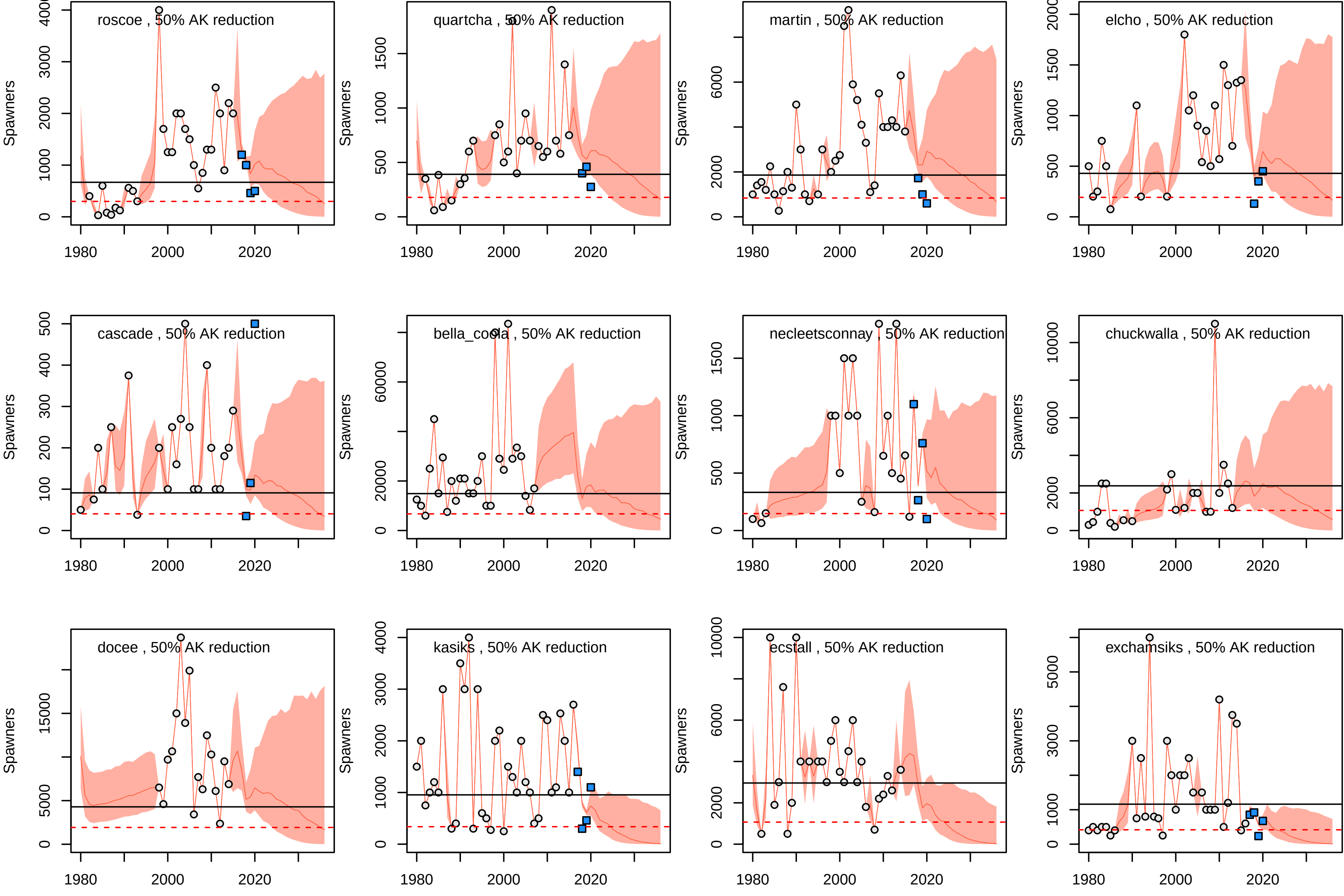
roscoe , 50% AK reduction
quartcha , 50% AK reduction
martin , 50% AK reduction
elcho , 50% AK reduction
cascade , 50% AK reduction
bella_coola , 50% AK reduction
necleetsconnay , 50% AK reduction
chuckwalla , 50% AK reduction
docee , 50% AK reduction
kasiks , 50% AK reduction
ecstall , 50% AK reduction
exchamsiks , 50% AK reduction
Spawners
1980
2000
2020

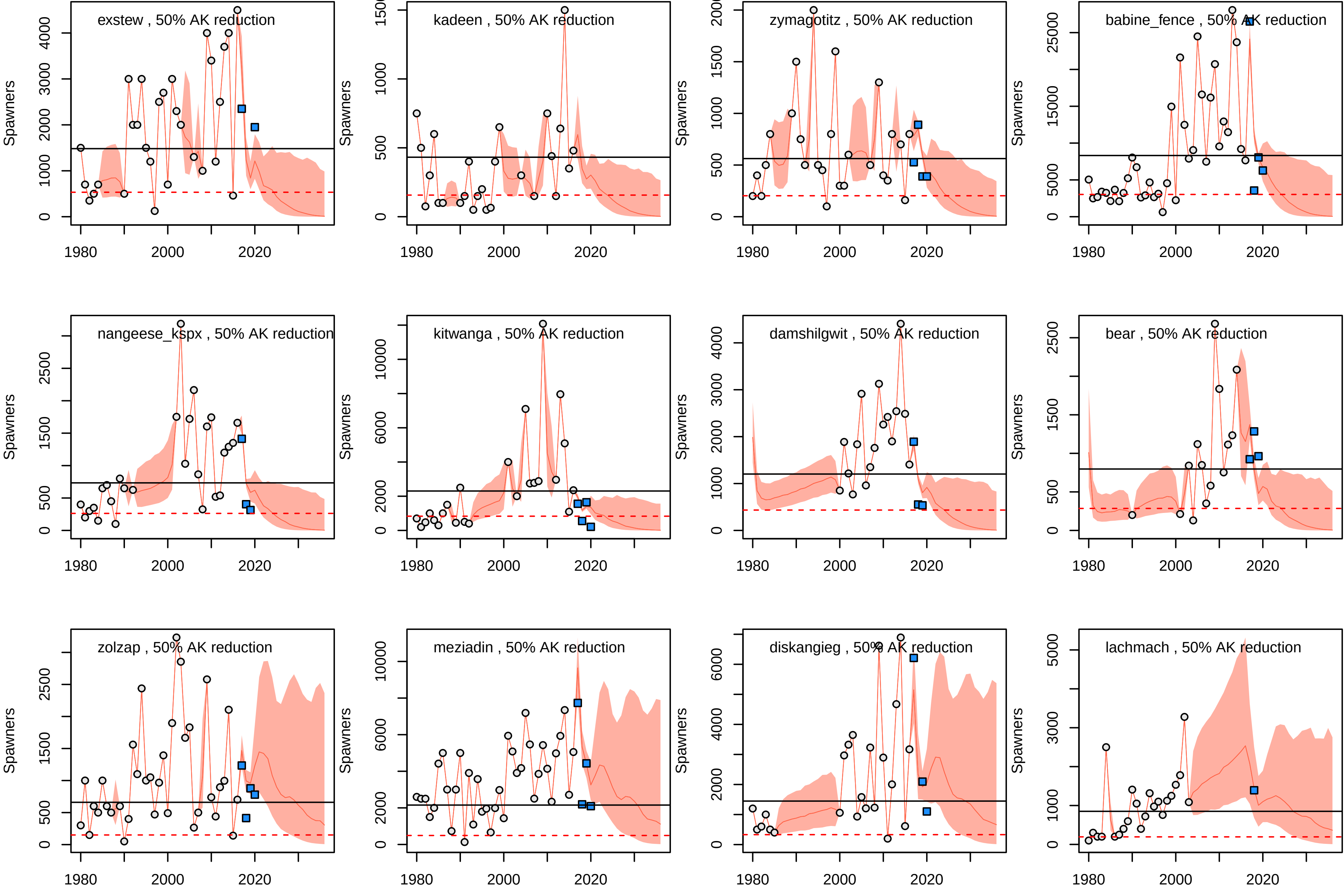
exstew , 50% AK reduction
kadeen , 50% AK reduction
zymagotitz , 50% AK reduction
babine_fence , 50% AK reduction
nangeese_kspx , 50% AK reduction
kitwanga , 50% AK reduction
damshilgwit , 50% AK reduction
bear , 50% AK reduction
zolzap , 50% AK reduction
meziadin , 50% AK reduction
diskangieg , 50% AK reduction
lachmach , 50% AK reduction
Spawners
1980
2000
2020

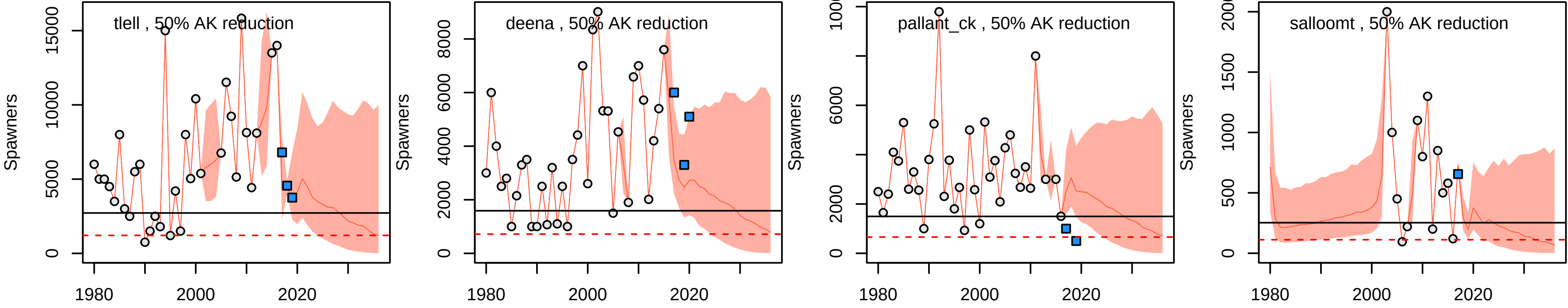

tlell , 50% AK reduction
deena , 50% AK reduction
pallant_ck , 50% AK reduction
salloomt , 50% AK reduction
Spawners
1980
2000
2020

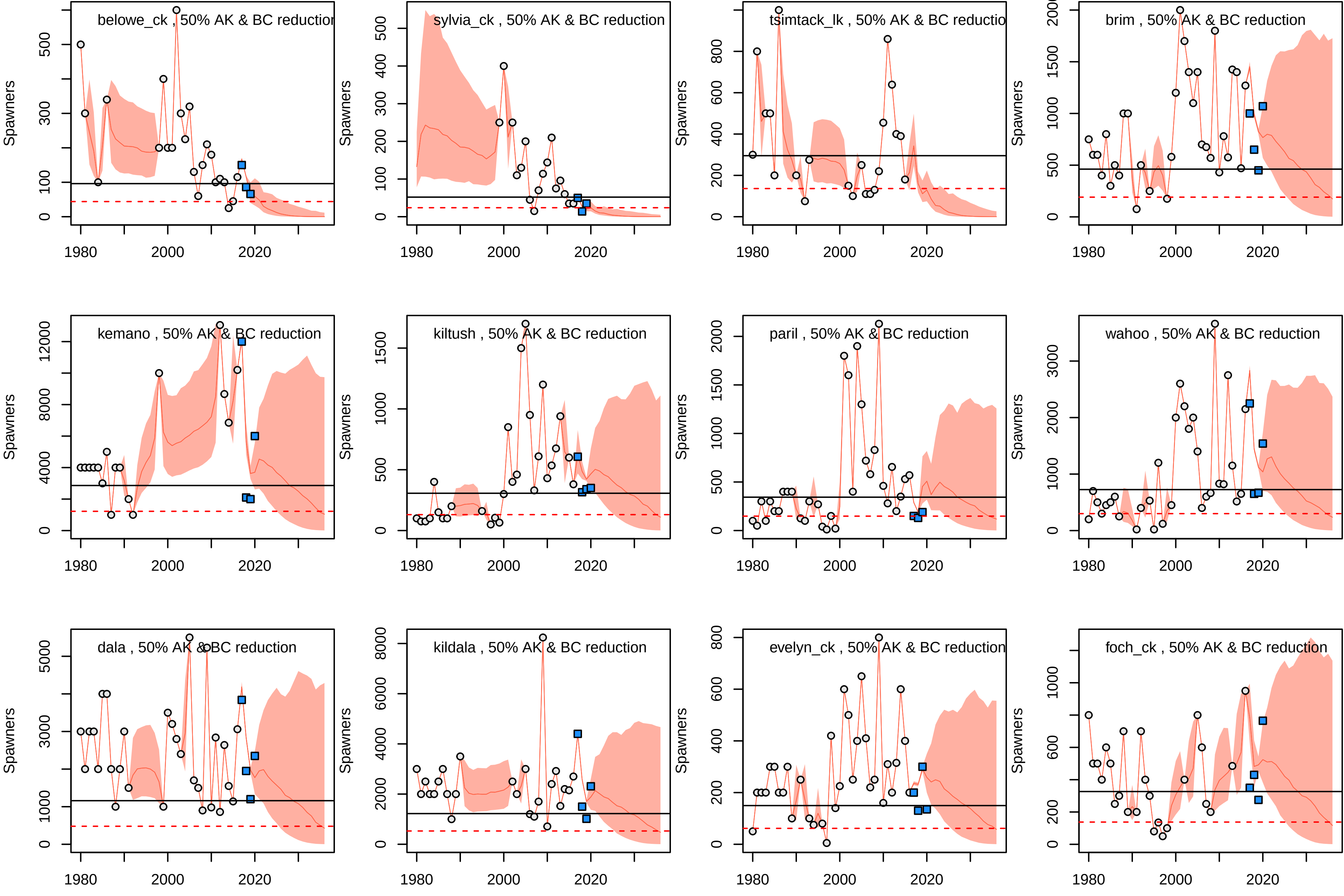
belowe_ck , 50% AK & BC reductio
sylvia_ck , 50% AK & BC reduction
tsimtack_lk , 50% AK & BC reductio
brim , 50% AK & BC reduction
kemano , 50% AK & BC reduction
kiltush , 50% AK & BC reduction
paril , 50% AK & BC reduction
wahoo , 50% AK & BC reduction
dala , 50% AK & BC reduction
kildala , 50% AK & BC reduction
evelyn_ck , 50% AK & BC reduction
foch_ck , 50% AK & BC reduction
Spawners
1980
2000
2020

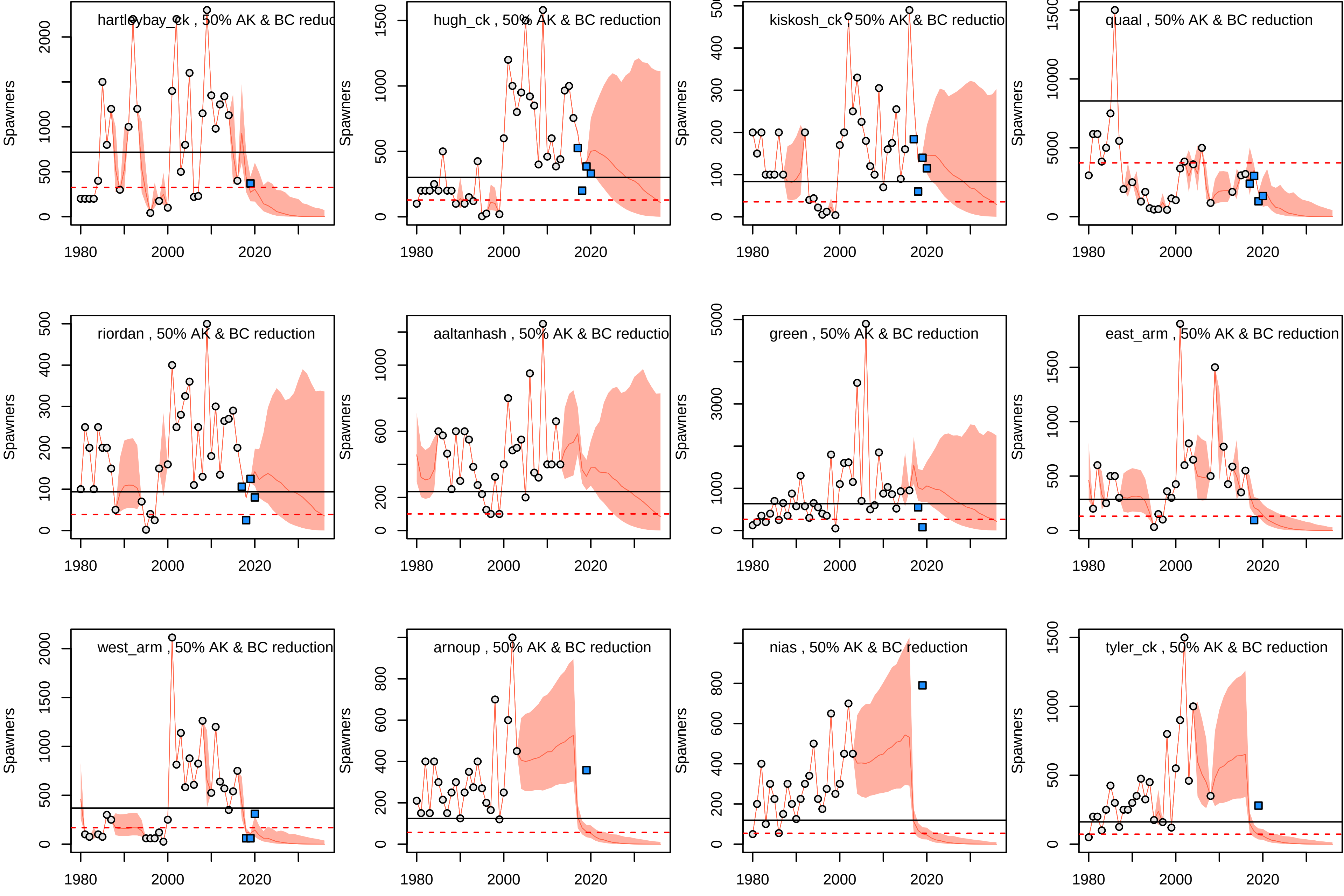
hartleybay_ck , 50% AK & BC reduc
hugh_ck , 50% AK & BC reduction
kiskosh_ck , 50% AK & BC reductio
quaal , 50% AK & BC reduction
riordan , 50% AK & BC reduction
aaltanhash , 50% AK & BC reductio
green , 50% AK & BC reduction
east_arm , 50% AK & BC reduction
west_arm , 50% AK & BC reduction
arnoup , 50% AK & BC reduction
nias , 50% AK & BC reduction
tyler_ck , 50% AK & BC reduction
Spawners
1980
2000
2020

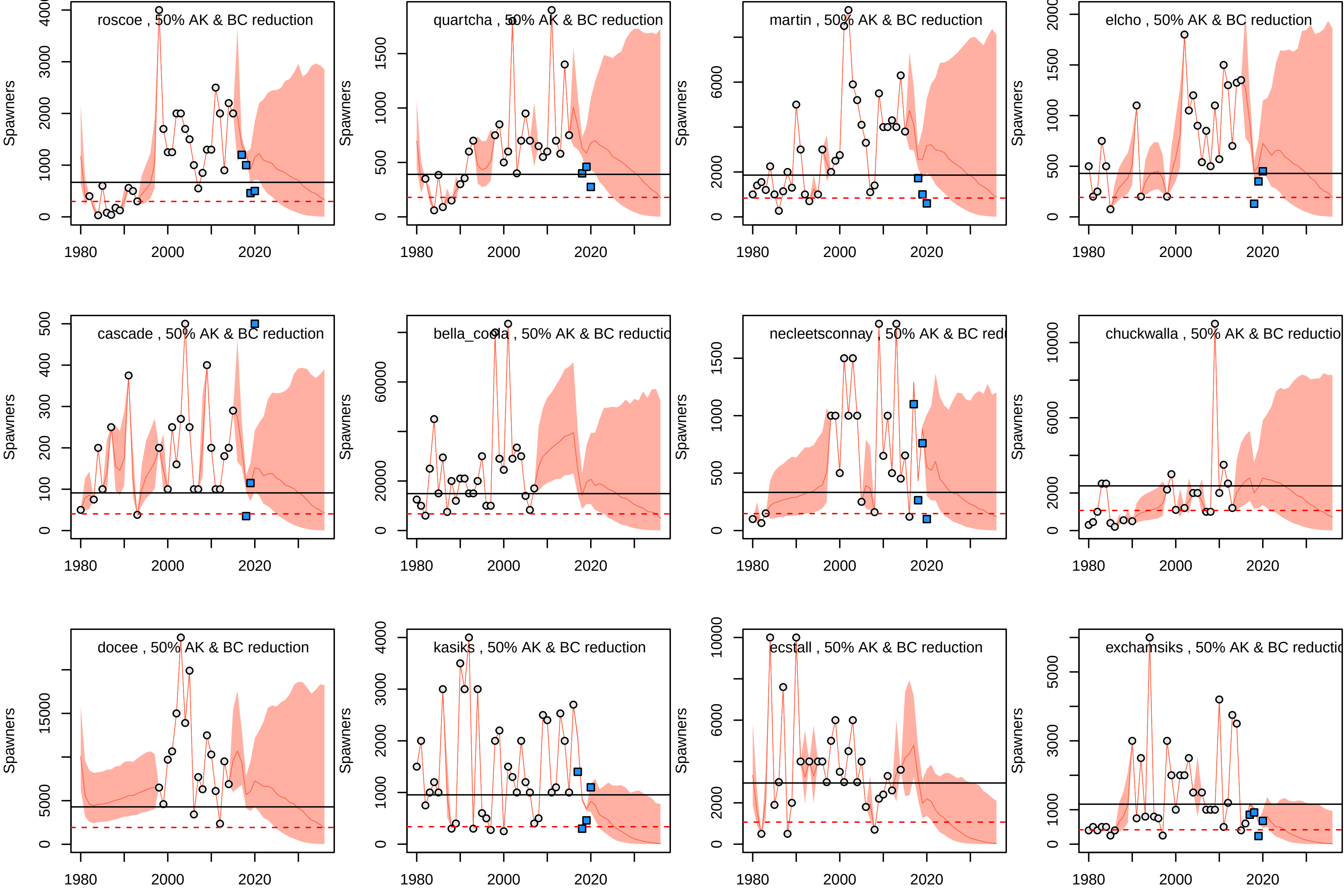

roscoe , 50% AK & BC reduction
quartcha , 50% AK & BC reduction
martin , 50% AK & BC reduction
elcho , 50% AK & BC reduction
cascade , 50% AK & BC reduction
bella_coola , 50% AK & BC reductio
necleetsconnay , 50% AK & BC redu
chuckwalla , 50% AK & BC reductio
docee , 50% AK & BC reduction
kasiks , 50% AK & BC reduction
ecstall , 50% AK & BC reduction
exchamsiks , 50% AK & BC reductio
Spawners
1980
2000
2020

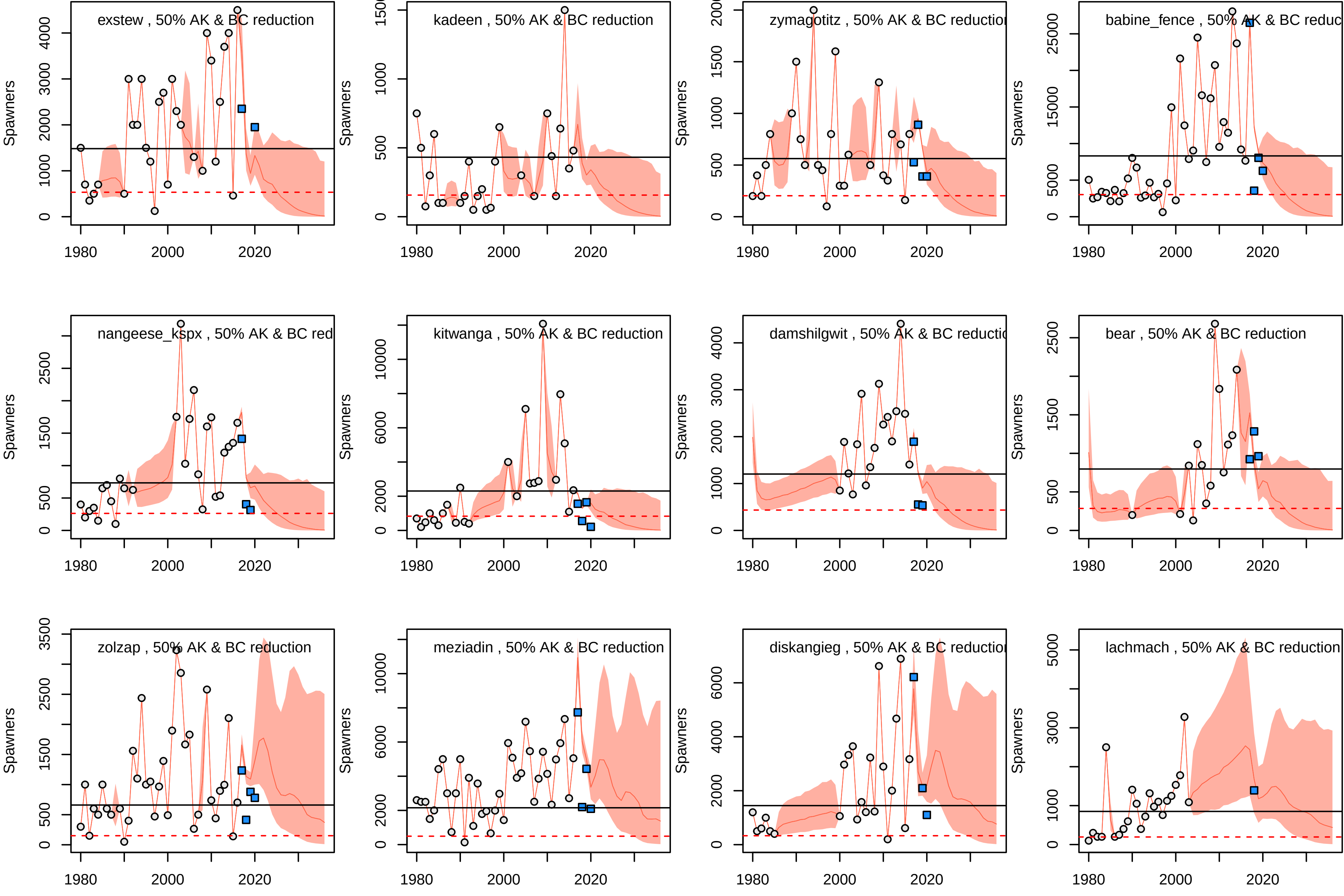
exstew , 50% AK & BC reduction
kadeen , 50% AK & BC reduction
zymagotitz , 50% AK & BC reductio
babine_fence , 50% AK & BC reduc
nangeese_kspx , 50% AK & BC red
kitwanga , 50% AK & BC reduction
damshilgwit , 50% AK & BC reductio
bear , 50% AK & BC reduction
zolzap , 50% AK & BC reduction
meziadin , 50% AK & BC reduction
diskangieg , 50% AK & BC reduction
lachmach , 50% AK & BC reduction
Spawners
1980
2000
2020

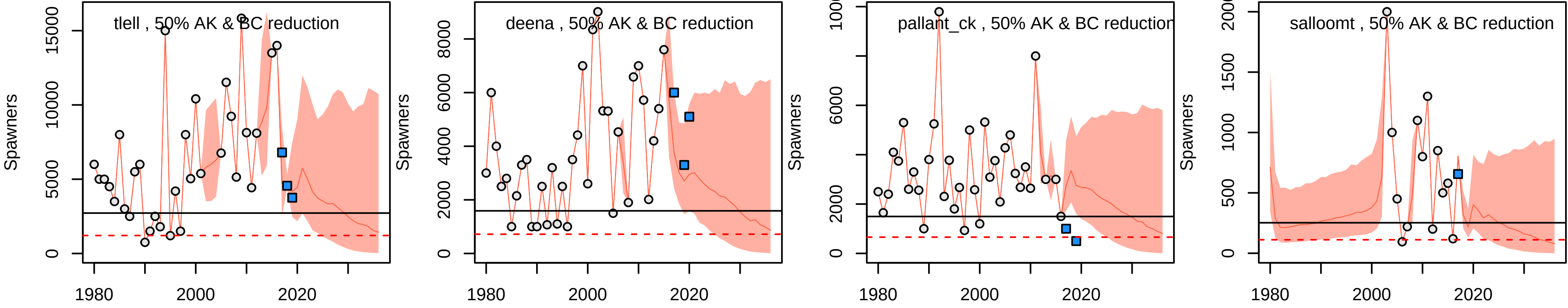

tlell , 50% AK & BC reduction
deena , 50% AK & BC reduction
pallant_ck , 50% AK & BC reduction
salloomt , 50% AK & BC reduction
Spawners
1980
2000
2020

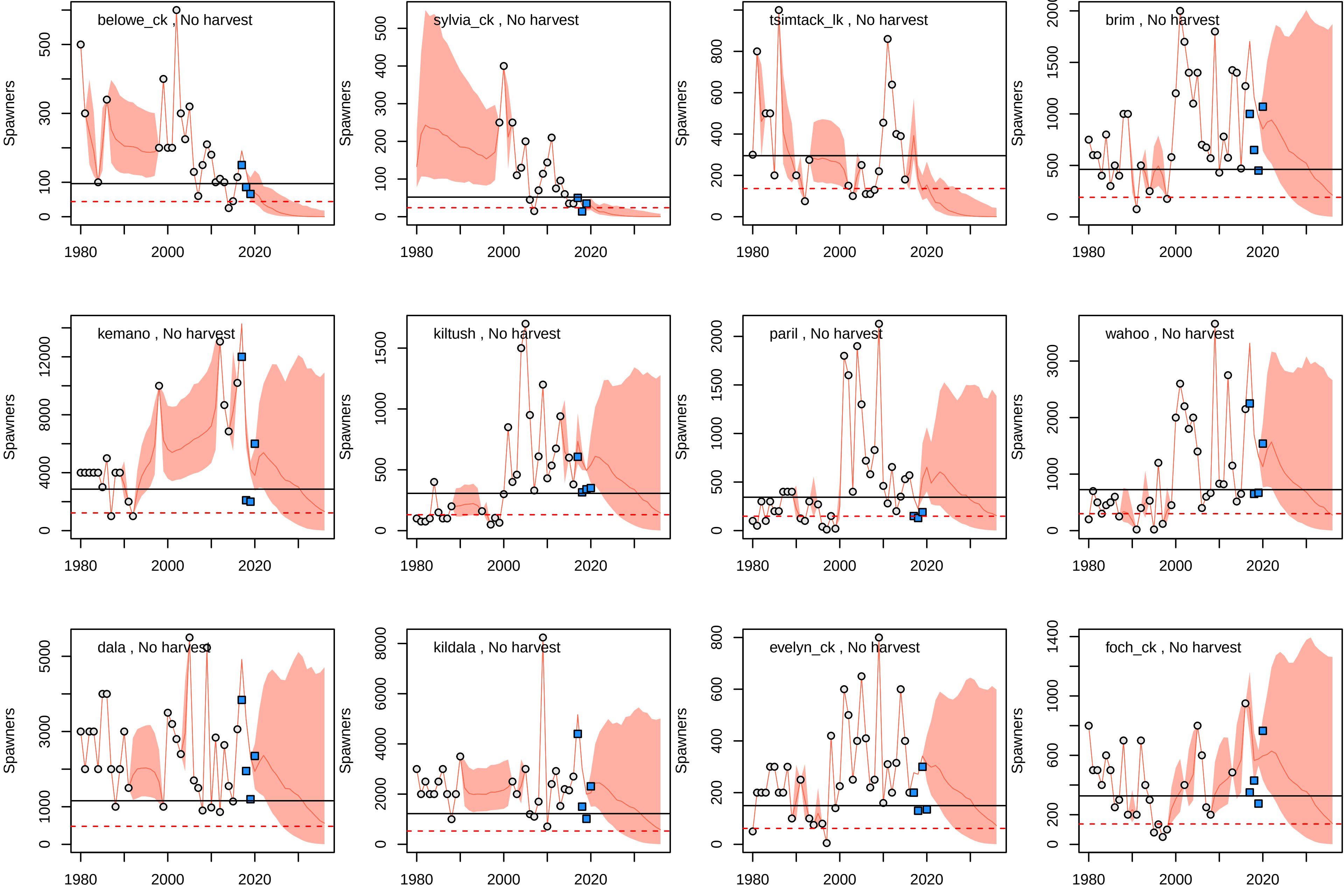

belowe_ck , No harvest
sylvia_ck , No harvest
tsimtack_lk , No harvest
brim , No harvest
kemano , No harvest
kiltush , No harvest
paril , No harvest
wahoo , No harvest
dala , No harvest
kildala , No harvest
evelyn_ck , No harvest
foch_ck , No harvest
Spawners
1980
2000
2020

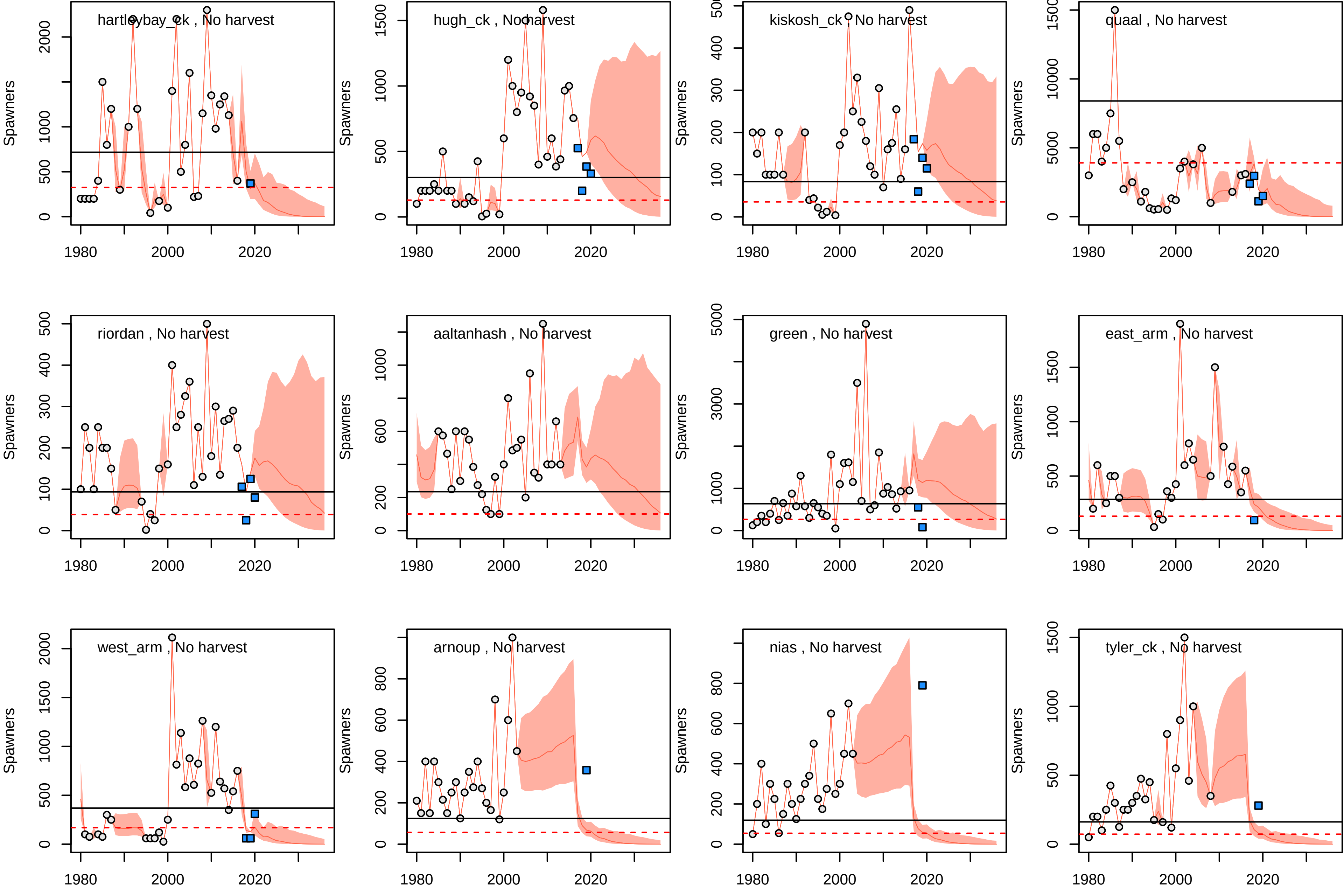
hartleybay_ck , No harvest
hugh_ck , No harvest
kiskosh_ck , No harvest
quaal , No harvest
riordan , No harvest
aaltanhash , No harvest
green , No harvest
east_arm , No harvest
west_arm , No harvest
arnoup , No harvest
nias , No harvest
tyler_ck , No harvest
Spawners
1980
2000
2020

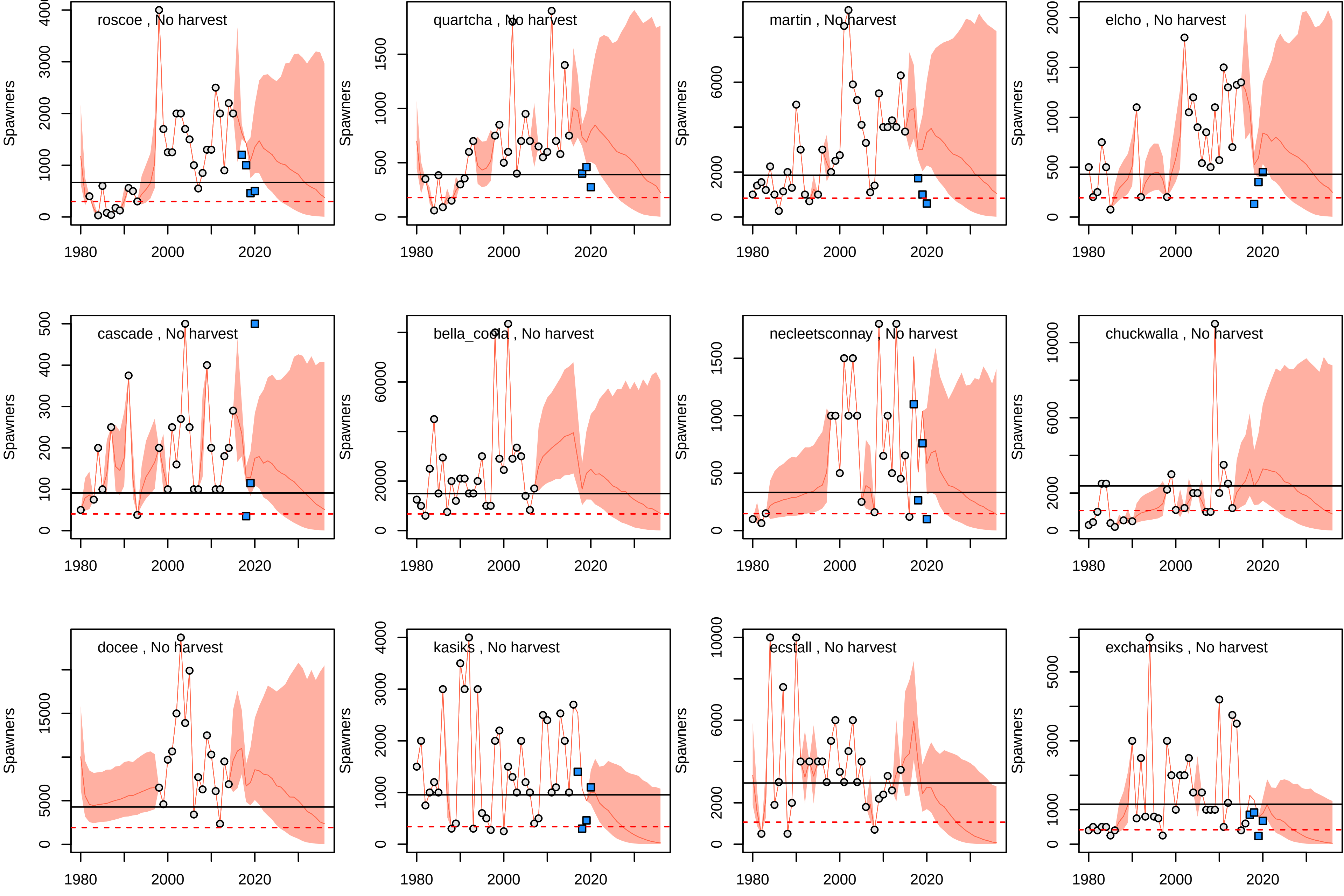
roscoe , No harvest
quartcha , No harvest
martin , No harvest
elcho , No harvest
cascade , No harvest
bella_coola , No harvest
necleetsconnay , No harvest
chuckwalla , No harvest
docee , No harvest
kasiks , No harvest
ecstall , No harvest
exchamsiks , No harvest
Spawners
1980
2000
2020

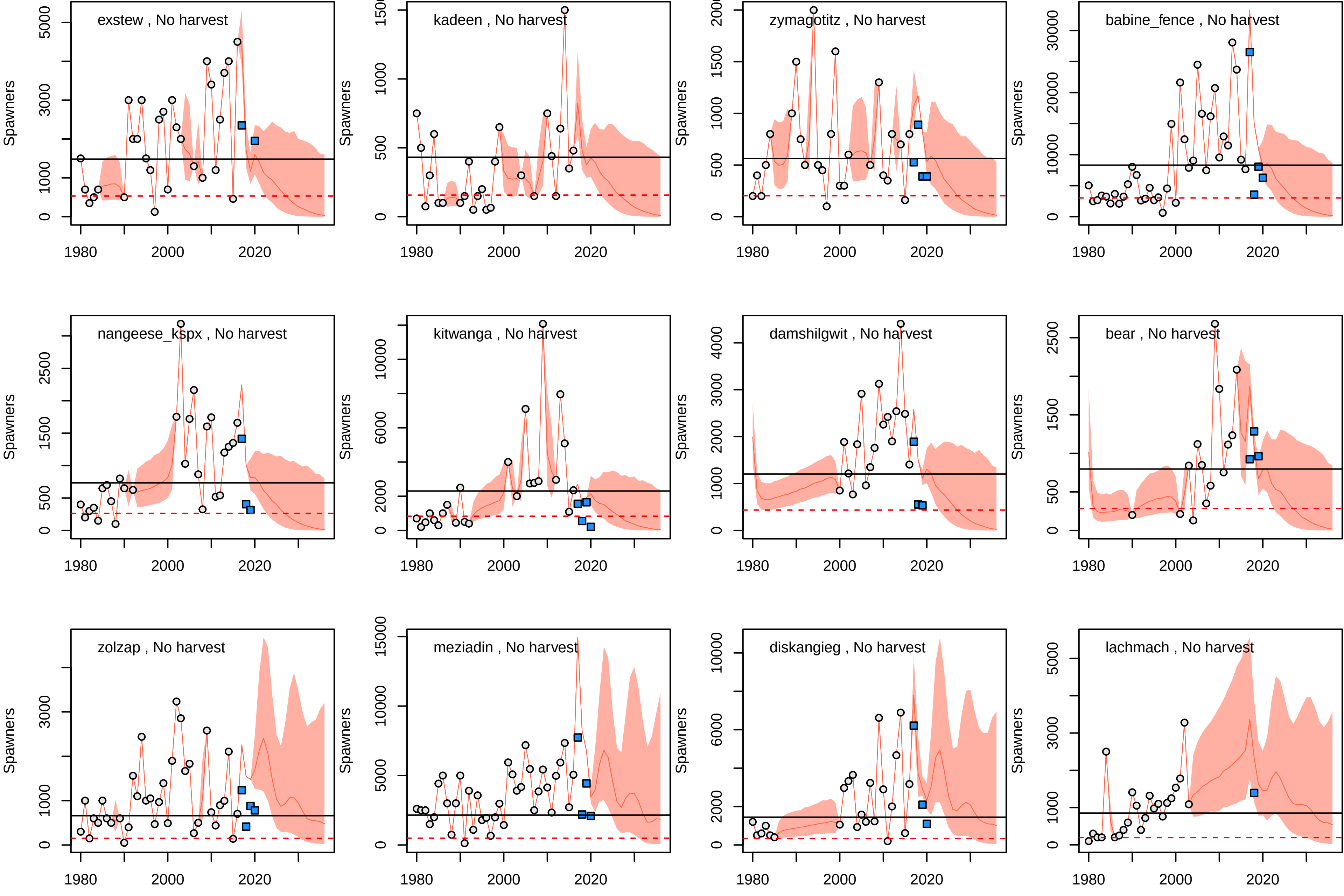
exstew , No harvest
kadeen , No harvest
zymagotitz , No harvest
babine_fence , No harvest
nangeese_kspx , No harvest
kitwanga , No harvest
damshilgwit , No harvest
bear , No harvest
zolzap , No harvest
meziadin , No harvest
diskangieg , No harvest
lachmach , No harvest
Spawners
1980
2000
2020

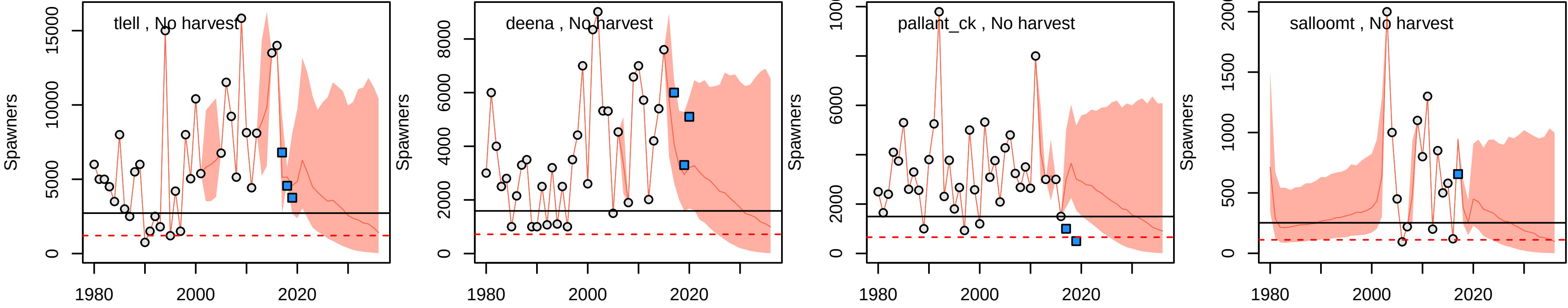
tlell , No harvest
deena , No harvest
pallant_ck , No harvest
salloomt , No harvest
Spawners
Spawners
Spawners
Spawners
0
5000
10000
15000
0
2000
4000
6000
8000
0
2000
6000
0
500
1000
1500
1980
2000
2020
1980
2000
2020
1980
2000
2020
1980
2000
2020
